# ABCA7 deficiency exacerbates glutamate excitotoxicity in Alzheimer’s disease mice – a new pharmacological target for Glu-related neurotoxicity

**DOI:** 10.1101/2025.07.25.666774

**Authors:** Anna Maria Górska, Irene Santos-García, Aleš Kvasnička, Dana Dobešová, David Friedecký, Jacob Gildenblat, Jens Pahnke

## Abstract

Increasing attention has been directed towards the perturbation of glutamate (Glu) and γ-aminobutyric acid (GABA) homeostasis during the pathogenesis of Alzheimer’s disease (AD). The prevailing disequilibrium, stemming from hyperactivation of the glutamatergic system, culminates in progressive neuronal impairment and cognitive deterioration. This study aimed to elucidate the contributory role of the ATP-binding cassette transporter A7 (ABCA7), identified as the second most critical genetic determinant in AD, in glutamatergic-associated neurotoxicity. This endeavor sought to advance molecular comprehension of neurological disorders where Glu-GABA neurotransmission represents a pivotal pharmacotherapeutic target.

Utilizing multi-omics approaches, we rigorously analyzed four distinct mouse models, both with and without APPtg and ABCA7 expression, to simulate varied pathological and ABCA7-deficient states. Our results revealed amyloid-beta (Aβ) deposition as a catalyst for surging glutamatergic transmission. Notably, ABCA7 ablation exacerbated glutamatergic-induced neurotoxicity, attributed to diminished enzymatic activity related to neurotransmitter degradation and amplified expression levels of specific neurotransmitter transport proteins and receptor subunits, notably NMDA, AMPA, and GABA_A_.

These findings furnish the first comprehensive description elucidating ABCA7’s amplification of neurotoxic effects through modulation of Glu-GABA neurotransmission systems in neurodegenerative contexts, primarily mediated by lipid interaction. The evidence underscores ABCA7’s imperative role in shaping future pharmacological strategies aimed at counteracting neurodegeneration precipitated by Glu-mediated neurotoxicity. This research advances the frontier for therapeutic exploration to ameliorate the deleterious neural consequences characteristic of neurodegenerative pathologies.

**Highlights:**

1. Alterations within the ABCA7 transporter locus constitute the second most significant genetic predisposition factor for Alzheimer’s disease (AD), subsequent to the influence of the APOE4 allele.
2. Excessive stimulation of glutamatergic neurotransmission culminates in excitotoxicity, leading to the gradual demise of neuronal populations due to pathological hyperactivity.
3. In murine models with wild-type genetics, the absence of ABCA7 results in diminished functionality of both the glutamatergic and GABAergic neurotransmitter systems.
4. Conversely, in mouse models engineered to mimic Alzheimer’s pathology, deficiency in ABCA7 exacerbates glutamate-induced neurotoxicity.
5. During amyloid-β accumulation, the absence of ABCA7 correlates with an elevation in specific lipid levels, potentially contributing to neurodegenerative processes.
6. From a therapeutic standpoint, pharmacological activation of ABCA7 may mitigate the neuronal death associated with glutamate overactivation in individuals afflicted by neurodegenerative disorders.

**Graphical abstract:** 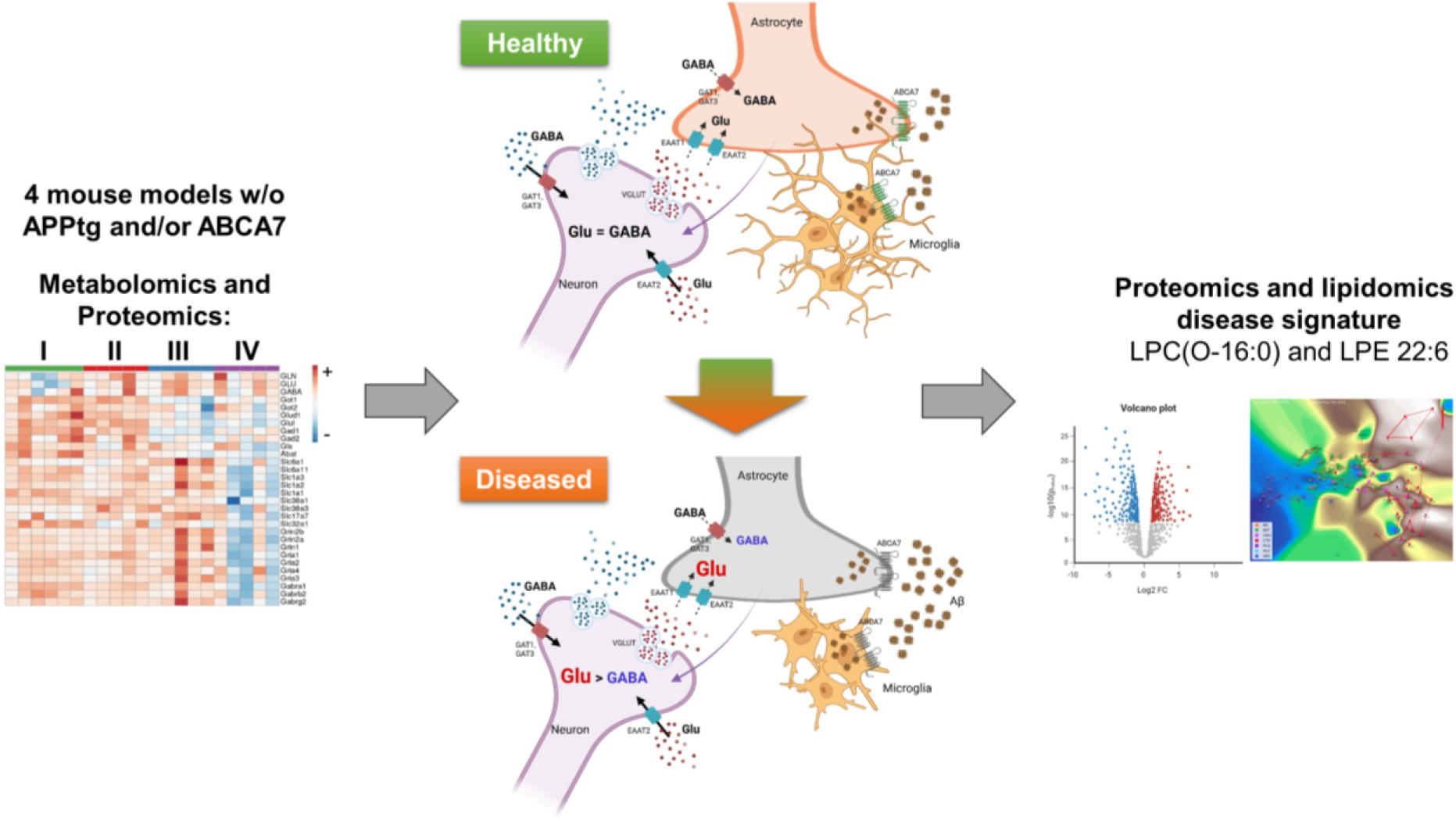

## 1 Introduction

Alzheimer’s disease (AD) ranks as the fifth leading cause of mortality among individuals aged 65 years and older and holds the position of the sixth leading cause of death for the adult population at large ^1^. In 2019, over 55 million individuals globally were affected by AD and other forms of dementia. According to estimations of the World Health Organization (WHO), this number is anticipated to escalate to approximately 140 million by the year 2050 ^1^. AD is characterized by the deposition of amyloid-β (Aβ) plaques and tau neurofibrillary tangles within the neocortex. However, the biochemical and cellular alterations occurring within the brain remain only partially elucidated ^2,3^. The most profound symptoms of AD include memory loss and disorientation, which are associated with a reduction in neuronal count within the hippocampus, a brain region fundamental to learning and memory ^4^. The entorhinal cortex, the interface between the hippocampus and the neocortex, is essential for forming spatial memory and is the initial region impacted by AD pathology ^5^.

In recent years, the significant correlation between the imbalance of amino acid neurotransmitters, specifically glutamate (Glu) and γ-aminobutyric acid (GABA), and the pathogenesis of AD has garnered considerable attention ^6^. Notably, the overstimulation of Glu coupled with insufficient inhibition by GABA is associated with progressive deposition of Aβ ^7,8^.

Glu, recognized as the predominant excitatory neurotransmitter within the mammalian central nervous system (CNS) and a precursor to GABA ^9^, is essential to processes such as memory, neuronal development, and synaptic plasticity. Its distribution is widespread throughout the CNS, predominantly localized within cortical and hippocampal pyramidal neurons that are integral to cognitive function ^10^. Most glutamatergic neurotransmission in the mammalian CNS is facilitated by ionotropic glutamate receptors (iGluRs) – specifically, NMDA and AMPA receptors. These receptors are pivotal in modulating synaptic plasticity and strength, thereby elucidating the molecular mechanisms underpinning learning and memory, which positions iGluRs as significant targets for therapeutic interventions ^11,12^. As of now, the only used drug mitigating the excessive stimulation of the glutamatergic system is memantine, a low-affinity antagonist of NMDA receptors that inhibits Glu-related neurotoxicity without disrupting the physiological functions of Glu that are necessary for memory and learning ^13^. The gradual decline in glutamatergic terminals may elucidate why memantine exhibits diminished efficacy in AD patients after several years of treatment.

Consequently, exploring potential new therapeutic strategies utilizing existing pharmacological agents that can counteract the progressive neurodegeneration and memory impairments associated with Glu overstimulation may provide substantial benefits to affected individuals. Currently, medications that influence the glutamatergic system are used in various neurological conditions such as:

- Riluzole in epilepsy ^14^ and amyotrophic lateral sclerosis ^15,16^, already used in frontotemporal dementia, is a neuroprotector that can decrease presynaptic Glu release by persistently blocking of calcium and sodium currents;
- Gabapentin, Pregabalin, and Perampanel ^17^ in epilepsy reduce Glu release;
- Topiramate in epilepsy, with AMPA antagonists’ properties, is also effective in patients with treatment-resistant schizophrenia ^18^;
- Fluoxetine (SSRI) and Imipramine (tricyclic antidepressant) in depression can restore AMPAR-mediated synaptic transmission ^19,20^;
- Ketamine in depression, an NMDAR blocker, is effective in treatment-resistant depression ^21^.

In 2010, a genome-wide association study (GWAS) identified the ATP-binding cassette, subfamily A, member 7 locus (*ABCA7*) as a genetic risk factor for AD ^22^. Since then, evidence derived from *in vitro*, *in vivo*, and human studies has substantiated the role of *ABCA7* as one of the most significant risk genes associated with AD ^23,24^. Our research, along with that of others, has demonstrated a robust correlation between the loss of ABCA7 function and the pathogenesis of AD ^25^. Patients exhibiting low levels of ABCA7 are at an increased risk of developing early-onset AD compared to those with the highest levels of ABCA7, which suggests that ABCA7 plays a protective role against the early stages of the disease, particularly through its function in the elimination of toxic lipids from cellular membranes ^26^. Furthermore, loss-of-function mutations in ABCA7 are associated with an increased risk of AD by 80 % in populations of African ancestry, and there is a reported increase of 100−400 % in the risk of early-onset AD in European ancestry populations ^23,27–29^. ABCA7 is widely distributed throughout the brain, with studies demonstrating its expression in neurons, astrocytes, microglia, endothelial cells of the blood–brain barrier (BBB), and brain pericytes in both rodents and humans ^30–33^. Under healthy conditions, ABCA7 is instrumental in maintaining lipid homeostasis in the brain and, significantly, facilitates the phagocytosis of Aβ aggregates by microglia ^34–36^. ABCA7, by its role in the cholesterol homeostasis ^33^ and phospholipids transport, represents one of the lipid-related risk factors associated with AD ^27,37^. The principal lipid classes known to be affected in AD encompass cholesterol, sphingolipids, phospholipids, and glycerolipids, including gangliosides ^38^. The human brain is abundant in lipids, and variances in their concentrations and distributions yield significant insights. The innovation of highly sensitive mass spectrometry and mass spectrometry imaging has enabled the identification of intricate alterations throughout the lipidome and their correlations with various disease states, thereby assisting in the identification of potential biomarkers ^39–42^.

In this work, we elucidated the disruption of Glu-GABA equilibrium in a murine AD model through the application of multi-omics analyses, emphasizing the significant role of ABCA7 in modulating Glu-mediated neurotoxicity associated with AD. Furthermore, by employing advanced methodologies ^42–44^, we are the first to identify lipids that appear specific to the pathology of AD concerning ABCA7 deficiency.

## 2 Materials and methods

### 2.1 Glu-Gln-GABA cycle markers for metabolomics and proteomics analyses

**Tab. 1.**
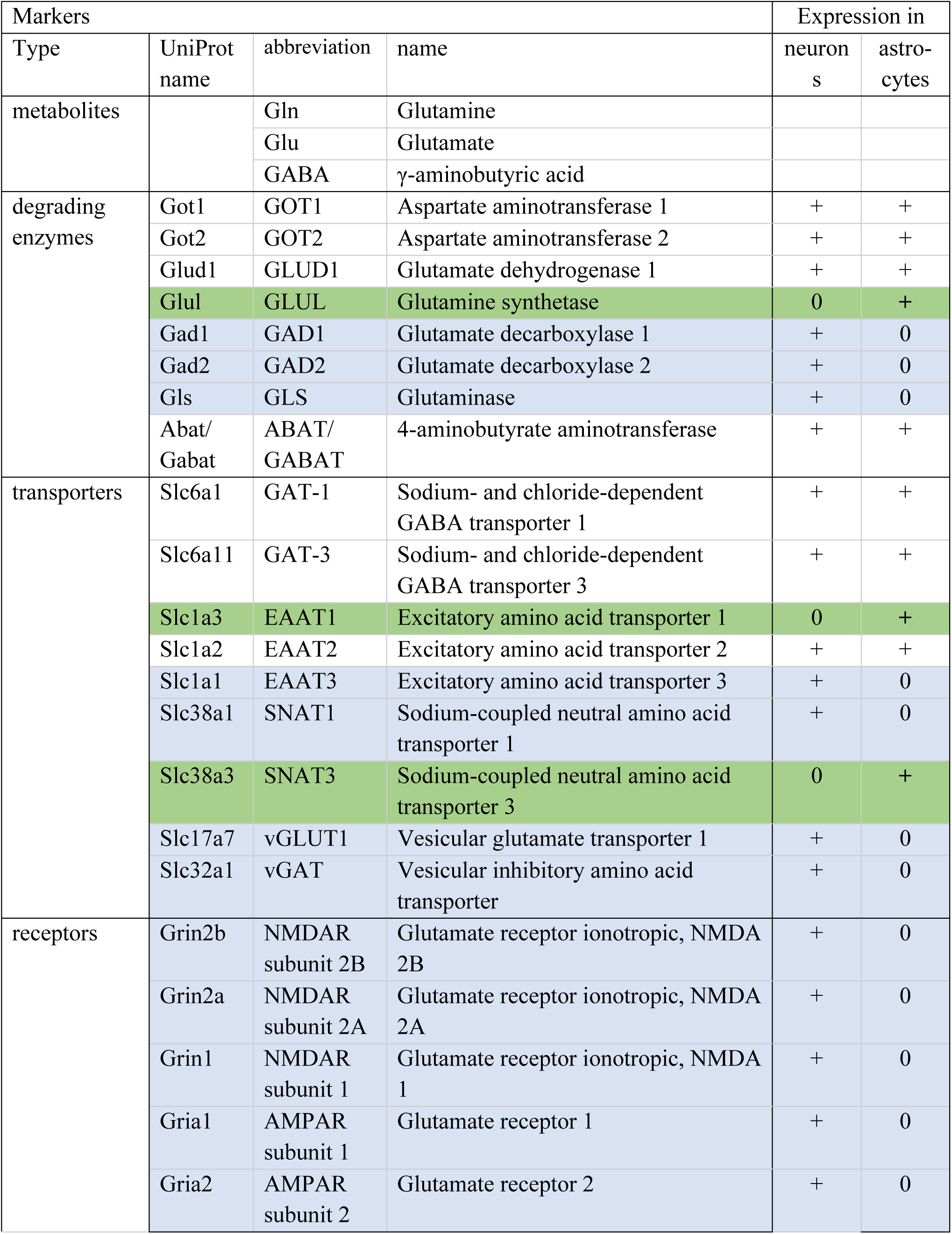
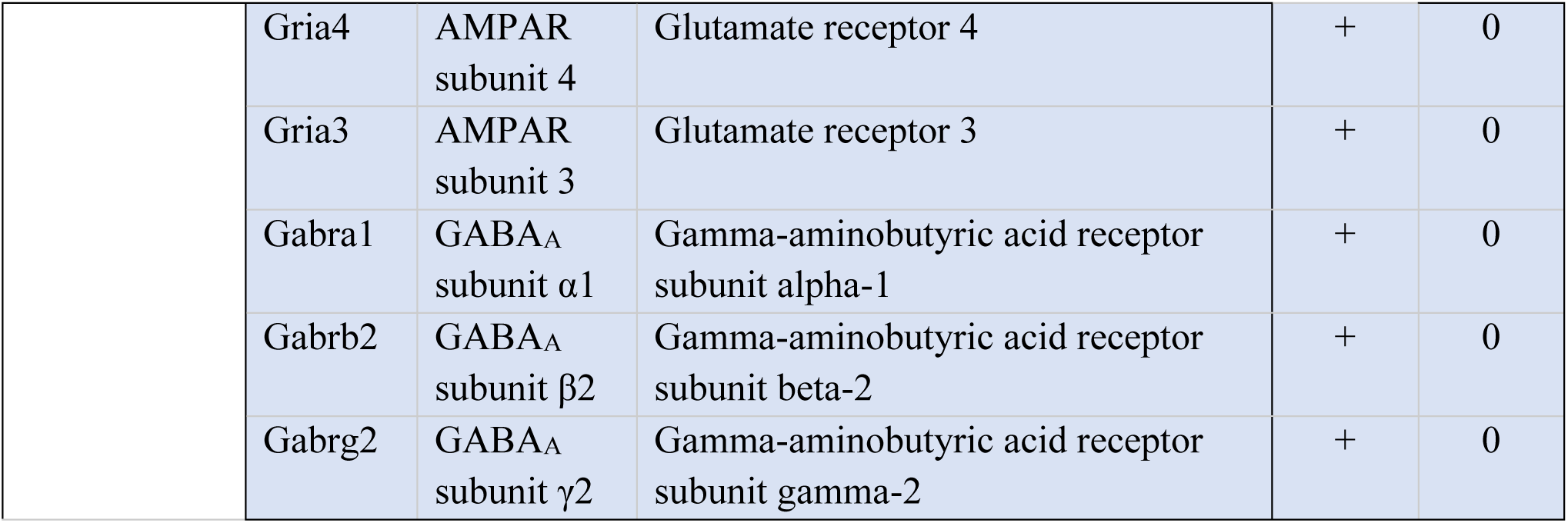
Analyzed markers of the Glu-Gln-GABA system (see also **Figures 1 and 2**). The markers are categorized into metabolites (Glu, Gln, and GABA) and proteins, including degrading enzymes, transporters, and receptors, with their expression in neurons and/or astrocytes. Three of the proteins are only expressed in astrocytes (green), 17 are specifically expressed in neurons (light blue).

### 2.2 Animal models and breeding scheme

Four distinct lines of mice, all in the C57BL/6J background, were compared in this study:

1. **APPtg** – APP/PS1 mice (B6.Cg-TgThy1-APPSw, Thy1-PSEN1*L166P/21^JckrPahnk^) ^45^,
2. **A7ko** – constitutive knockouts of the Abca7tm1.1^(*ABCA7*)Pahnk^ allele (generated as described in ^46^),
3. **APP-A7ko** – APPPS1 mice (number 1) crossed with Abca7tm1.1^(***ABCA7***)Pahnk^ knockouts (number 2), and
4. **WT** – C57BL/6J wild-type littermates.

In all experimental animals, the APP/PS1 transgene was present in hemizygous form (APPtg: +/0), and the *Abca7* knockout was homozygous (−/−). The experimental animals were generated by crossing hemizygous (APPtg: +/0) and heterozygous (*Abca7*: +/−) sires with *Abca7* heterozygous (+/−) dams. The resulting offspring were allocated to the different experimental groups according to their genotypes.

The animals were housed at the Department of Comparative Medicine (section Radium Hospital) at Oslo University Hospital (Norway) under the following macroenvironmental conditions: temperature: 22 ± 1 °C; relative humidity: 62 ± 5%; ventilation: 15 air changes per hour; light cycle: 12 hours dark/light; illuminance: 1 lux at night, 70 lux during the day, 400 lux for working illumination. The animals were placed in Eurostandard type III cages (Makrolon®), with groups of up to 8 individuals per cage. Aspen wood (*Populus tremula*, Tapvei®, Estonia) served as a bedding substrate and was changed once per week. Additional enrichment materials (tissue paper, tunnel rods, and occasionally gnawing sticks) were also provided in all the cages. Mice were offered food *ad libitum* (maintenance expanded pellets from SDS, Estonia) and acidified water at pH 3. Health monitoring was performed three times per year according to FELASA guidelines, and opportunistic organisms were included in one of the tests. Only segmented filamentous bacteria were detected during the studies, and they were not considered relevant or a cause of bias in the results. All experiments were conducted following the guidelines for animal experimentation by the European Union and Norwegian national laws.

### 2.3 Tissue collection and processing

Mice were euthanized by cervical dislocation. After transcardial perfusion with ice-cold PBS, brains were removed. Tissue was snap-frozen in liquid nitrogen and stored at −80°C until LC-MS/MS analysis.

### 2.4 Collection of cerebrospinal fluid (CSF) and interstitial fluid (ISF)

CSF and ISF samples were collected from WT (n=6), APPtg (n=6), APP-A7ko (n=5–6), and A7ko (n=6) mice that were 50 and 200 days old, respectively, following the protocol ^47^. Approximately 4 µl of CSF from the lateral ventricle (LV) were collected per mouse. Collected CSF was immediately transferred into a 100µl reaction tube, snap-frozen in liquid nitrogen, and stored at −80 °C until further analysis. The obtained CSF samples were analyzed individually (not pooled) later. In a separate group of animals, microdialysis probes (AT12.8.1.PE, 3 MDa cut-off, AgnTho’s AB, Sweden) for ISF collection were implanted perpendicular to the cortex surface (coordinates relative to bregma: a-p, +0.4 mm; m-l, ±3.0 mm; d-v, −2.2 mm, according to ^48^. Microdialysis probes were perfused with artificial cerebrospinal fluid (aCSF) [mM]: 147 NaCl, 2.7 KCl, 1.0 MgCl_2_, 1.2 CaCl_2_, pH=7.4) at a flow rate of 0.5 μl/min using a syringe pump (U-801, 8301300, AgnTho’s AB, Sweden). ISF was collected on ice for 1 h, then snap frozen in liquid nitrogen and stored at −80 °C until further LC-MS/MS analysis.

### 2.5 Proteomics analyses

#### 2.5.1 Brain tissue sample preparation

Brain tissue samples from WT (n=5–6), APPtg (n=5), APP-A7ko (n=3–5), and A7ko (n=3–5) mice aged 50, 100, 150, and 200 days were used as biological replicates for proteomics analyses. For each replicate, equal amounts of brain homogenate (approximately 20 µg of protein) were precipitated on amine beads, as previously described ^49^. The precipitated proteins on the beads were dissolved in 50 mM ammonium bicarbonate, reduced, alkylated, and digested with trypsin (1:50 enzyme:protein ratio; Promega, USA) at 37 °C overnight. The digested peptides were acidified and desalted on EVOTIPs using a standard protocol from EVOSEP (EVOSEP Biosystems, Denmark).

#### 2.5.2 LC-MS/MS proteomics analysis of brain tissue samples

Liquid chromatography with tandem mass spectrometry (LC-MS/MS) analysis was performed using an EVOSEP one LC system coupled with a timsTOF pro2™ mass spectrometer and a CaptiveSpray nanoelectrospray ion source (Bruker Daltonics GmbH, Bremen, Germany). 200 ng of the digested peptides were loaded onto a capillary C18 column (15 cm length, 150 μm inner diameter, 1.5 μm particle size, EVOSEP, Odense, Denmark). Peptides were separated at 50 °C using EVOSEP’s standard 30 sample/day method. The timsTOF pro2™ mass spectrometer operated in data-dependent Parallel Accumulation-Serial Fragmentation (PASEF®) mode ^50^. Mass spectra for MS and MS/MS scans were recorded in the m/z range of 100 to 1,700 Da. Ion mobility resolution was set between 0.85 and 1.40 V·s/cm over a ramp time of 100 ms. Data-dependent acquisition was conducted using four PASEF MS/MS scans per cycle with a nearly 100 % duty cycle. A polygon filter was applied in the m/z and ion mobility space to filter out low m/z, singly charged ions from PASEF precursor selection. An active exclusion time of 0.4 minutes was applied to precursors that reached 20,000 intensity units. The collisional energy was ramped stepwise as a function of ion mobility.

#### 2.5.3 Proteomics data processing of brain tissue samples

Raw data files from LC-MS/MS analyses were submitted to MaxQuant software (version 2.4.3.0, Max-Planck-Institute of Biochemistry, Martinsried, Germany) for protein identification and quantification ^51,52^. The UniProt mouse database (UniProt Consortium, European Bioinformatics Institute, EMBL-EBI, UK) was used. Trypsin without proline restriction enzyme option was used, with two allowed miscleavage sites. Carbamidomethyl was set as a fixed modification, and acetyl (protein N-term), carbamyl (N-term), and oxidation (M) were set as variable modifications. First, a search peptide tolerance of 20 ppm and a main search error of 4.5 ppm were used. The allowed false discovery rate (FDR) was 0.01 (1 %) for peptide and protein identification. Label-free quantitation (LFQ) was employed with default settings.

#### 2.5.4 CSF and ISF sample preparation

CSF and ISF samples from WT (n=6), APPtg (n=6), APP-A7ko (n=5-6), and A7ko (n=6) mice were used as biological replicates for the proteomics analyses. Samples were collected according to established protocol ^47^, for each replicate equal volumes of fluid (approx. 2 µg of protein) were precipitated on amine beads as previously described ^49^. The precipitated proteins on beads were dissolved in 50 mM ammonium bicarbonate, reduced, alkylated, and digested with trypsin (1:50 enzyme:protein ratio; Promega, USA) at 37 °C overnight. Digested peptides were acidified and desalted on EVOTIPs using a standard protocol from EVOSEP.

#### 2.5.5 LC-MS/MS proteomics analysis of CSF and ISF samples

LC-MS/MS analysis was carried out using an EVOSEP one LC system (EVOSEP Biosystems, Denmark) coupled to a timsTOF fleX™ mass spectrometer, using a CaptiveSpray nano electrospray ion source (Bruker Corporation, Germany). 200 ng of digested peptides were loaded onto a capillary C18 column (15 cm length, 75 μm inner diameter, 1.7 μm particle size, 120 Å pore size; IonOpticks, Fitzroy, VIC, Australia). Peptides were separated at 50 °C using the standard 40 sample/day method from EVOSEP. The timsTOF fleX™ mass spectrometer was operated in dia-PASEF® mode ^50^. Mass spectra for MS were recorded between m/z 100 and 1,700. Ion mobility resolution was set to 0.85–1.30 V·s/cm over a ramp time of 100 ms. The MS/MS mass range was limited to m/z 475– 1,000, and ion-mobility resolution to 0.85–1.27 V s/cm to exclude singly changed ions. The estimated cycle time was 0.95 s with eight cycles using DIA windows of 25 Da. Collisional energy was ramped from 20eV at 0.60 V s/cm to 59eV at 1.60 V s/cm.

#### 2.5.6 Proteomics data processing of CSF and ISF samples

Raw data files from LC-MS/MS analyses were submitted to data-independent acquisition proteomics data processing (DIA-NN, version 1.8.1) for protein identification and label-free quantification using the library-free function and the mobility module for dia-PASEF analysis ^53,54^. The UniProt mouse database (UniProt consortium, European Bioinformatics Institute, EMBL-EBI, UK) was digested for library generation. Carbamidomethyl (C) was set as a fixed modification. Trypsin without proline restriction enzyme option was used, with one allowed miscleavage, and peptide length range was set to 7–30 amino acids. The mass accuracy was set to 15 ppm, and precursor FDR allowed was 0.01 (1%).

### 2.6 Metabolomics analyses

#### 2.6.1 Brain tissue sample preparation

Brain tissue samples from WT (n=6), APPtg (n=6), APP-A7ko (n=4–6), and A7ko (n=3–6) mice aged 50, 100, 150, and 200 days were used as biological replicates for metabolomics analyses. Metabolites were extracted from brain homogenates using a protocol based on a previous study ^55^. Tissue samples (10 mg) were combined with 200 µL of methanol containing internal standards, mixed with water in a 4:1 (v/v) ratio, and vortexed for 10 s. The mixture was stored at –80 °C for 2 h for deproteinization. Following thawing, the samples were vortexed for another 10 s and centrifuged at 15,000 g for 15 min at 4 °C. The supernatant (approximately 160 µL) was transferred to LC-MS glass vials. A pooled quality control (QC) sample was created by combining 5 µL from each final extract.

#### 2.6.2 LC-MS/MS metabolomics analysis

Targeted metabolomics analysis was conducted using an ExionLC ultra-high-performance liquid chromatograph coupled with a QTrap 6500+ mass spectrometer (Sciex, Foster City, CA, USA). The setup was controlled using Analyst software (v1.6.2). A HILIC column (Luna 3 μm NH2, 2 × 100 mm, Phenomenex) was employed to separate metabolites. Mobile phase A consisted of 20 mM ammonium acetate (pH=9.75), while mobile phase B was acetonitrile. The column temperature was set at 35 °C, with a 0.3 mL/min flow rate. The gradient settings were as follows: 0 min: 95 % B; 7 min: 10 % B; 13 min: 10 % B; 13.5 min: 95 % B; and 17 min: 95 % B. The total runtime was 17 min. Polarity switching allowed for the acquisition of metabolites in both positive and negative ionization modes. Optimal declustering potentials and collision energies were established using reference standards. Detailed methodology for detecting 352 metabolites in biological samples has been reported previously ^56^.

#### 2.6.3 Metabolomics data processing

Data were processed using SCIEX OS software (v1.6.1) and the Metabol package in R (v4.0.3, https://github.com/AlzbetaG/Metabol/). QC-based, locally estimated smoothing signal (LOESS) correction was applied, and analytes with a coefficient of variation (CV) above 30 % in the QC samples were excluded. Internal standards were used to monitor sample preparation consistency and instrument stability. Data were further normalized to tissue weight, and the normalized peak areas were used to present metabolomic data.

### 2.7 Lipidomics analyses

#### 2.7.1 Brain tissue sample preparation

Brain tissue samples from WT (n=6), APPtg (n=6), APP-A7ko (n=4–6), and A7ko (n=3–6) mice aged 50, 100, 150, and 200 days were used as biological replicates for lipidomics analyses. Lipids were isolated from brain tissue homogenates using a modified two-phase extraction method based on Ding et al. ^57^. Briefly, samples were kept on ice throughout the preparation process. Methanol (225 µL) containing deuterated internal standards (3% SPLASH LIPIDOMIX, 5 µmol/L oleic acid-d9, and 5 µmol/L ceramide d18:1–d7/15:0) was added to 10 mg of lyophilized brain tissue. After vortexing for 10 s, 750 µL of methyl tert-butyl ether was added, vortexed again for 10 s, and shaken at 4 °C for 15 min. Water (188 µL) was added to induce phase separation, followed by vortexing for 20 s. The samples were centrifuged at 14,000 g for 10 min, and 400 µL of the upper non-polar layer was collected and lyophilized overnight. The dried lipids were reconstituted in 200 µL of a mixture of acetonitrile, isopropanol, and water (2:2:1 v/v/v) before transfer to LC-MS vials. The QC sample was prepared similarly to the metabolomics QC.

#### 2.7.2 LC-MS/MS lipidomics analysis

Lipidomic profiling was performed using an ExionLC ultra-high-performance liquid chromatograph interfaced with a QTrap 6500+ mass spectrometer (Sciex, Concord, CA, USA) as previously described ^58^. Lipids were separated on a BEH C8 reversed-phase column (2.1 mm × 100 mm, 1.7 µm, Waters), tempered at 55 °C with a flow rate of 0.35 mL/min. The mobile phase A (MPA) was acetonitrile/water (3:2, v/v), and the mobile phase B (MPB) was isopropanol/acetonitrile (9:1, v/v), both containing 10 mM ammonium acetate. The elution gradient was: 0–1.5 min: 32% MPB; 1.5–15.5 min: increase to 85% MPB; 15.6–18 min: 97% MPB; 18.1–20 min: return to 32% MPB. Polarity switching allowed the simultaneous detection of lipids in positive and negative modes. QC samples were analyzed every sixth injection to ensure system stability.

#### 2.7.3 Lipidomics data processing

Data were processed using SCIEX OS software (v1.6.1) and the Metabol package in R (v4.0.3, https://github.com/AlzbetaG/Metabol/). QC-based, locally estimated smoothing signal (LOESS) correction was applied, and analytes with a coefficient of variation (CV) above 30 % in the QC samples were excluded. Internal standards were used to monitor sample preparation consistency and instrument stability. Data were further normalized to tissue weight, and the normalized peak areas were used to present lipidomic data.

### 2.8 Behavioral assessment - measurement of locomotor activity

Spontaneous locomotor activity in mice was measured every 10 min for 3 days (72 h) using an infrared ray passive sensor system (TSE PhenoMaster Systems, Germany). An apparatus with an infrared beam sensor was positioned on top of a conventional polypropylene cage, where multiple movements were counted and relayed to a computer interface. Mice aged 50 days (n=9–10) and 200 days (n=4–7) were utilized for the experiment. They were housed in standard cages and maintained on a 12-hour light/12-hour dark cycle with light onset at 06:00 a.m. Data were analyzed with Prism (v9, Dotmatics, Boston, USA) using a two-tailed unpaired Student’s t-test after performing a normality check for statistical significance between groups.

### 2.9 Statistical analyses

LFQ proteomic data were analyzed using Perseus (v2.0.11, https://cox-labs.github.io/coxdocs/perseus_instructions.html) ^59^. Multi-omics data were visualized with the Heat map and Principal component (PCA) analyses using ClustVis: a web tool for visualizing clustering of multivariate data (https://biit.cs.ut.ee/clustvis/) ^60^. Volcano plots presenting up- or down-regulated lipids between genotypes were generated using VolcaNoseR (https://huygens.science.uva.nl/VolcaNoseR/) ^61^, showing the effect size (fold of change, log_2_ transformation) vs. significance (*p* value, -log_10_ transformation). The data points with the effect size ±1.5 (up- or down-regulation) and a significance threshold > 1.3 (–lg0.05, corresponding to p < 0.05) were considered hits. All the statistical analyses were performed using GraphPad Prism software (v9, Dotmatics, Boston, MA, USA), after normality check using Shapiro‒Wilk normality test. If the data pass the normality check, two-tailed unpaired Student’s t-test was performed to determine the significant differences between the groups. The differences were considered significant when p < 0.05. The data are presented as the mean ± standard deviations (SD).

## 3 Results

### 3.1 Descriptive characterization of the OMICs datasets

#### 3.1.1 Proteomics analysis

Using a proteomics approach, we detected 7636 proteins in whole brain tissue samples, and 5961 and 788 proteins in CSF and ISF, respectively. Four groups of animals were selected to describe the importance of ABCA7 for the pathogenesis of AD: WT controls, APPtg, APP-A7ko, and A7ko (non-AD control). Further details are found in the Methods section.

In brain tissue, all markers of the Glu-Gln-GABA system were detected, as well as in CSF samples (**Table 1**). Moreover, in brain tissue, the following most relevant, directly AD-related proteins were detected: amyloid-beta precursor protein (APP), apolipoprotein E (APOE), presenilin-1 (PSEN1), and nicastrin (NCSTN). We also decided to analyze the level of the glial fibrillary acidic protein (GFAP), a marker associated with astrogliosis and linked to pathological processes in AD, particularly Aβ build-up and neuroinflammation ^62^. APP and GFAP were also detected in ISF samples, representing the extracellular pool of proteins.

#### 3.1.2 Metabolomics analysis

Using a metabolomics approach, 145 metabolites were detected in the brain tissue of WT, APPtg, APP-A7ko, and A7ko mice at 50, 100, 150, and 200 days of age. Three crucial metabolites, Glu, Gln, and GABA, were selected to support our proteomics data.

#### 3.1.3 Lipidomics analysis

Using the lipidomics approach, 112 lipids were detected in the whole brain tissue from WT, APPtg, APP-A7ko, and A7ko mice at 50, 100, 150, and 200 days of age. In our analysis, lipids from the following listed classes were detected: FA – Fatty acids, PG – Phosphatidylglycerols, PI – Phosphatidylinositols, PS – Phosphatidylserines, Cer – Ceramides, Hex2Cer – Dihexosylceramides, LPC – Lysophosphatidylcholines, and LPE – Lysophosphatidylethanolamines. The analyzed lipids from each class are listed in **Supplementary Material Fig. S14.**

### 3.2 ABCA7 deficiency leads to exacerbation of the Glu-Gln-GABA system of APPtg mice

Heat map analysis revealed that most of the changes were observed regarding neurotransmitter-degrading enzymes, transporters, and receptors (Fig. 1A, C, ovals and rectangles). Moreover, principal component analysis (PCA) highlights differences between control and APPtg mice as compared to the ABCA7 knockout groups (APP-A7ko and A7ko) (Fig. 1B, D). For better visualization of the differences between genotypes during the disease, the levels of degrading enzymes (Fig. S3A, C), transporters, and receptors (Fig. S3B, D) were presented using PCA, comparing early and late disease stages at 50 and 200 days of age. Based on the heat maps, the differences were predominantly observed in the APP-A7ko and A7ko groups. Thus, the variability of degrading enzymes (Fig. S3E, G), transporters, and receptors (Fig. S3F, H) between the samples was present during the disease progression from 50 days through 200 days of age.

**Fig. 1.**
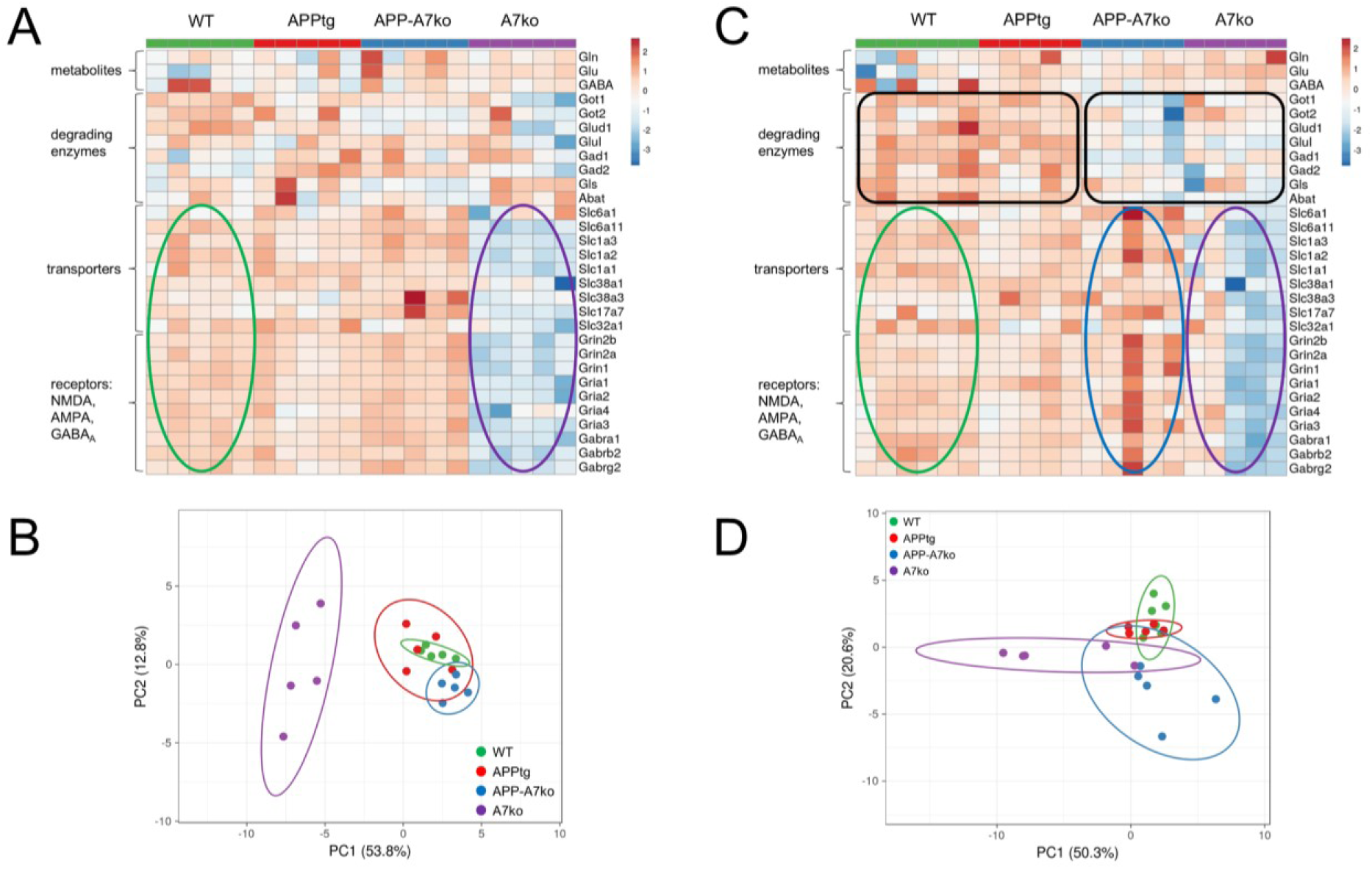
The lack of ABCA7 increases the activity of the Glu-Gln-GABA cycle in APPtg mice while decreasing it in WT controls. Heat map analysis of metabolites (Glu, Gln, GABA) and proteins involved in the Glu-Gln-GABA turnover and neurotransmission of 50-day-old (A) and 200-day-old (C) mice [background controls (WT), APPtg, APP-A7ko, and disease controls (A7ko)]. Plot of the principal component analysis (PCA) showing the differences between samples due to the genotype effect in (B) 50 and (D) 200-day-old mice.

In 50-day-old animals, changes were only observed in A7ko animals (Fig. 1B), where downregulation of the transporters and all examined receptors for Glu and GABA was detected (Fig. 1A shows expression; in Fig. 1B, it separates from other models). This was analyzed using a two-tailed unpaired Student’s t-test, GAT1 and SNAT1 p>0.05 (n.s.). At the age of 200 days, changes occurred in animals with accumulations of Aβ and/or ABCA7 transporter dysfunction (APP-A7ko and A7ko).

APP-A7ko mice showed a significant decrease in degrading enzymes expression at 50 days of age (values are given as mean[lg(intensity)] ± SD): 17.96±1.36 vs 18.07±1.35 (APPtg) and 18.04±1.50 (WT); at 200 days: 17.85±1.53 vs 18.25±1.46 (APPtg) and 18.29±1.42 (WT); two-tailed unpaired Student’s t-test, 200 days, APP-A7ko vs. B6 and APPtg; GLUL, GLS, GOT2 p>0.05, n.s.)) which was concomitant with an increase in transporter (GAT1, EAAT2, SNAT1) and receptor subunit (GRIN2B, GRIA3) levels (50 days: 16.78±2.00 vs 16.52±1.88 (APPtg) and 16.58±1.96 (WT); 200 days: 16.82±2.09 vs 16.64±1.91 (APPtg) and 16.71±1.96 (WT); two-tailed unpaired Student’s t-test, 200 days, APP-A7ko vs. APPtg)) (Fig. 1C, black rectangle and blue oval; Fig. S3C and D).

Moreover, in these animals, disease progression (50 vs. 200 days of age) primarily affects the activity of degrading enzymes and the levels of transporters and receptors (Fig. S3E and F).

However, the most surprising result was the effect of the A7ko on the Glu-Gln-GABA cycle. In A7ko mice, at 50 days of age, marked downregulation (fold of change WT vs. A7ko = 0.44; APP-A7ko vs. A7ko = 0.39) of transporters and receptors was observed (mean [lg(intensity)] ± SD for 50 days: 15.42±2.03 (A7ko) vs 16.77±2.00 (APP-A7ko), 16.58±1.96 (WT); two-tailed unpaired Student’s t-test; GAT1 and SNAT1 p>0.05, n.s.) (Fig. 1A, purple oval; and 1B, Fig. S3B), followed by downregulation of degrading enzymes (WT vs. A7ko fold of change = 0.75; WT vs. APP-A7ko = 0.74) (200 days: 17.88±1.40 (A7ko) / 17.85±1.53 (APP-A7ko) vs. 18.29±1.42 (WT); two-tailed unpaired Student’s t-test; GOT2 and GAD2, p>0.05, n.s.)) in late age (200 days of age) (Fig. 1C, purple oval; Fig. S3G and H).

### 3.3 Dysregulation of Glu and GABA metabolism *at the astrocytic and neuronal levels* is induced by Aβ deposition and lack of ABCA7

The balance between the metabolism of Glu and GABA is maintained by mutual communication between astrocytes and neurons. Here, astrocytes remove excessive Glu and GABA from the synaptic cleft and then supply neurons with glutamine (Gln) for Glu and GABA synthesis ^63^. Gln is transported to neurons by SNAT1 and SNAT3 ^64–66^ and then metabolized into Glu in glutamatergic neurons and GABA in GABAergic neurons *via* glutaminase (GLS) ^67–69^, and glutamate decarboxylase (GAD), respectively (Fig.2). After the release of neurotransmitters into the synaptic cleft, the excess of Glu and GABA is captured by neighboring astrocytes using specific transporters, excitatory amino acid transporters (EAAT1 and 2) and sodium- and chloride-dependent GABA transporters (GAT-1 and -2). Highly toxic Glu is immediately broken down *via* glutamine synthetase (GLUL) into Gln ^63^. At the same time, GABA is metabolized by GABA transaminase (GABA-T, ABAT) to succinic semi-aldehyde (SSA), followed by succinate, which is incorporated into the tricarboxylic acid (TCA) cycle ^69^ (Fig. 2).

**Fig. 2.**
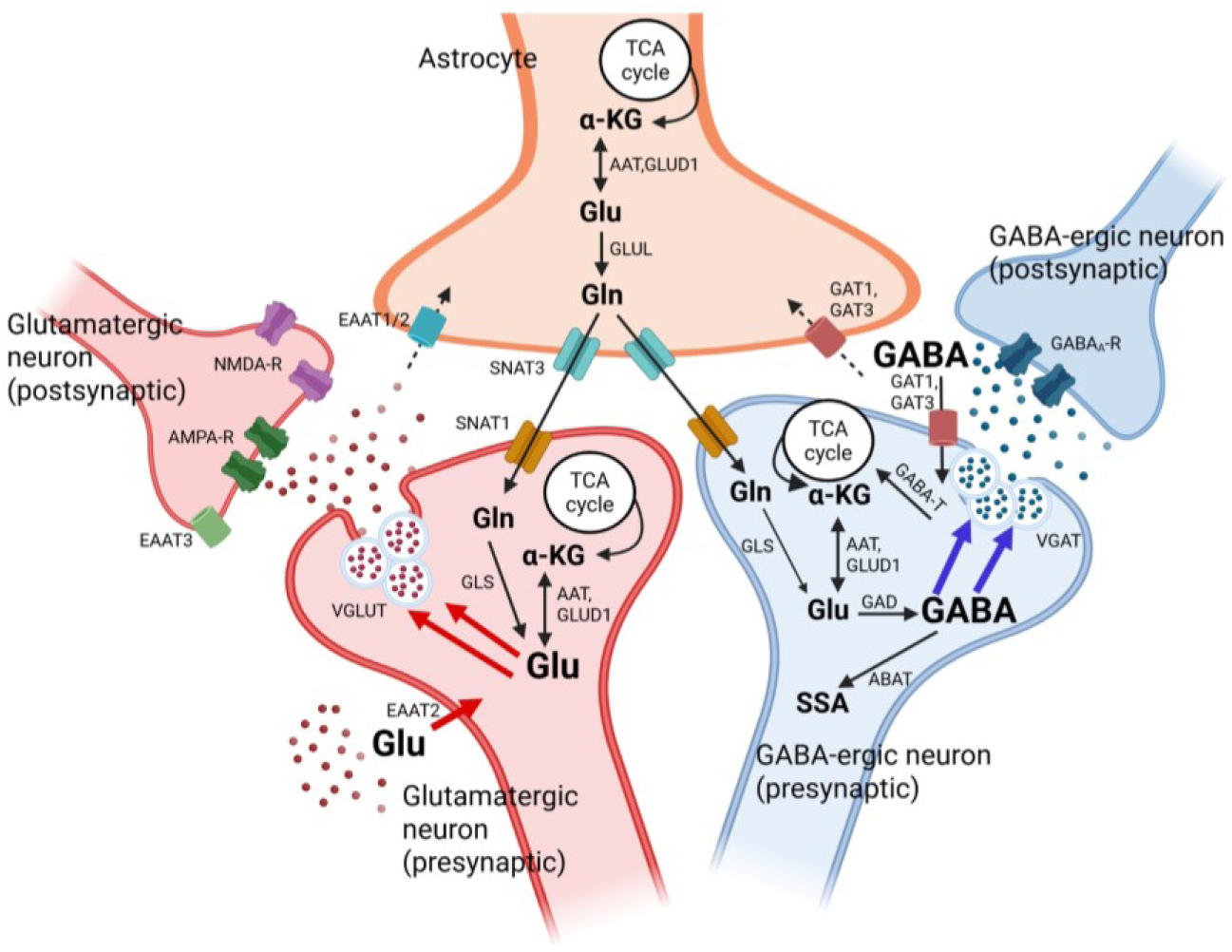
Glu-Gln-GABA cycle in pre- and postsynaptic neurons and astrocytes. The maintenance of these neurotransmitter pools strictly depends on the *de novo* synthesis of glutamine (Gln) in astrocytes, followed by the rapid and efficient removal of neurotransmitters from the synaptic cleft to maintain homeostasis and provide Gln for replenishing neurotransmitter pools in both glutamatergic and GABAergic neurons. α-KG, α-ketoglutarate; Glu, glutamate; Gln, glutamine; GABA, γ-aminobutyric acid; SSA, succinic semialdehyde; GABA-T/ABAT, 4-aminobutyrate aminotransferase; AAT, aspartate aminotransferase, GLUD1, glutamate dehydrogenase 1; GLUL, glutamine synthetase; GLS, glutaminase; GAD, glutamate decarboxylase; GAT1/3, Sodium- and chloride-dependent GABA transporter 1/3; SNAT1/3, Sodium-coupled neutral amino acid transporter 1/3; vGLUT, vesicular glutamate transporter; vGAT, vesicular inhibitory amino acid transporter; EAAT1/2/3, excitatory amino acid transporter 1/2/3; AMPA-R, AMPA receptor; NMDA-R, NMDA receptor; GABA_A_-R, GABA_A_ receptor.

An imbalance between inhibitory and excitatory neurotransmission is an essential feature of AD ^70^. Here, the level of Glu and GABA is measured in the whole brain, meaning the pool which is released into the synaptic cleft and stored inside the neuronal vesicles. Thus, changes in the level of degrading enzymes may also be affected by neurotransmitter pools inside the neurons and the efficiency of reuptake transporters. In physiological conditions, Glu and GABA are mainly metabolized by astrocytes instead of being taken up into neurons and packed again into vesicles.

Our results showed that in AD mice the lack of ABCA7 results in decreased levels of the enzymes (AAT/GOT1 and GLUD1) responsible for converting Glu into α-ketoglutarate (α-KG). Simultaneously, the stable GLUL levels suggest a reduction in Glu metabolism at the astrocytic level. Furthermore, the increased activity of EAAT2 drives Glu into astrocytes, where impaired Glu metabolism may lead to higher astrocytic toxicity and excess Glu in the synaptic cleft. In pathological conditions, Glu-related toxicity is intensified by impaired Aβ plaque clearance *via* ABCA7. Interestingly, the lack of ABCA7 in control animals (A7ko) impaired Glu and GABA metabolism at the astrocytic and neuronal levels without changing the neurotransmitter levels, suggesting that already-synthesized Glu and GABA are stored inside the vesicles.

GOT1 (AAT) and GLUD1 are reverse enzymes that convert αKG to Glu and Glu back into αKG (Fig. 3). These enzymes are expressed in both neurons and astrocytes, with higher activity observed in astrocytes, making them the primary cell type responsible for Glu metabolism. In APPtg animals, a decrease in GLUD1 was seen in 50- and 100-day-old animals (Fig. 4B), while no differences were found in GLUL activity (Fig. 4D). The changes in enzyme levels were accompanied by an increase in Glu (Fig. 5C) with no changes in Gln levels (Fig. 4F), suggesting that neuronal metabolism is favored in APPtg mice compared to astrocytes. This is supported by decreased GLS activity, which is specifically expressed in neurons. Additionally, impairment of Glu metabolism at the astrocytic level may also lead to a persistent excess of Glu in the synaptic cleft. In the GABAergic system, an increase in GAD2 (GAD65) (Fig. 4H) was observed in 50-day-old animals, with no changes in GAD1 (GAD67) (Fig. 4G), followed by no difference in GABA levels and a decrease in ABAT (Fig. 4I and J).

**Fig. 3.**
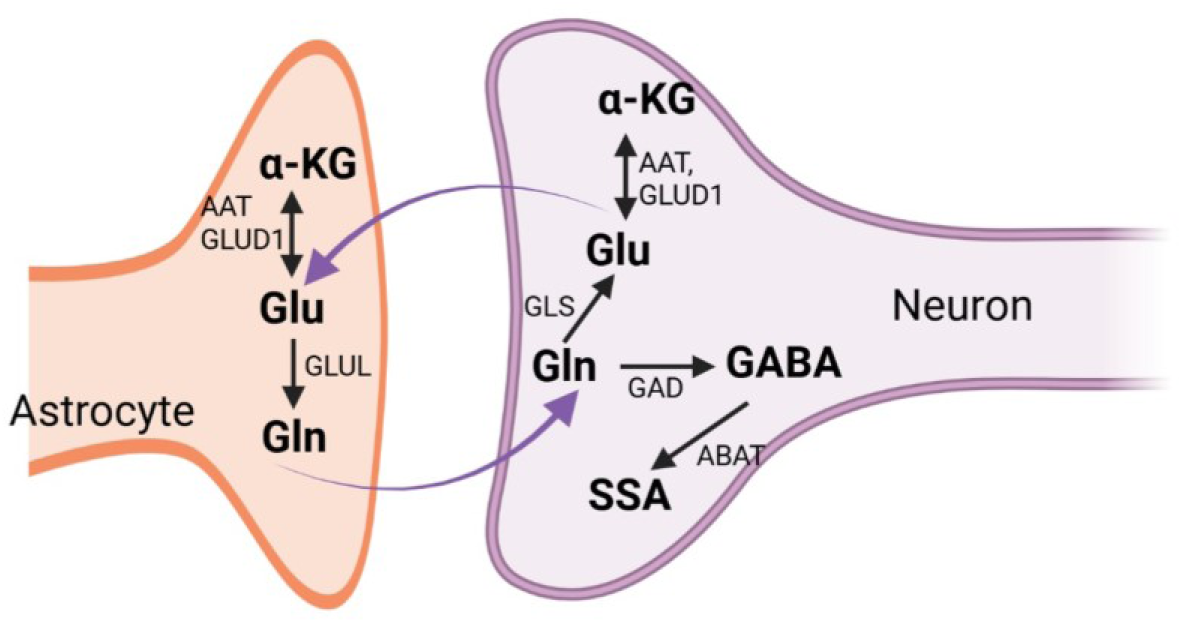
Astrocyte – neuron cycle of Glu-Gln-GABA synthesis and metabolism. Astrocytes provide neurons with Gln, a precursor for synthesizing Glu and GABA. Astrocytes absorb excess Glu to complete the cycle, which is then metabolized back into Gln.

**Fig. 4.**
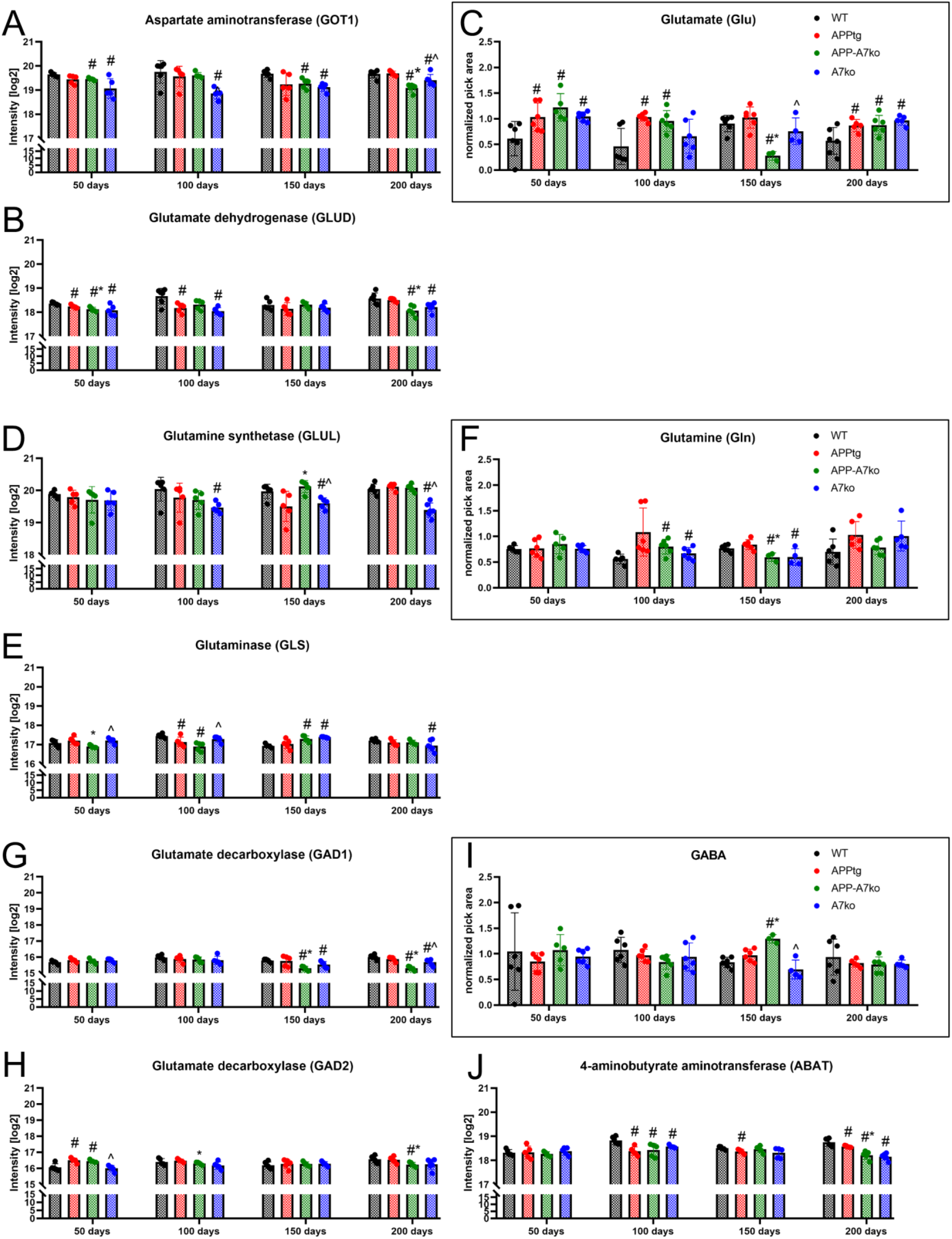
AD pathology favors neuronal reuptake, which persists Glu longer in the synaptic cleft. Lack of functional ABCA7 transporters in healthy/control animals inhibits Glu and GABA turnover in astrocytes and neurons. Diagram present the combined metabolomics (C) Glu, (F) Gln and (I) GABA (in black rectangular) and proteomics data for degrading enzymes: (A) GOT1, (B) GLUD, (D) GLUL, (E) GLS, (G) GAD1, (H) GAD2 and (J) ABAT. After a normality check, statistical analyses were performed with a two-tailed unpaired Student’s t-test. p < 0.05 was considered to be significant, * vs APPtg; ^ vs APP-A7ko; # vs WT. Data are presented as the mean ± SD.

**Fig. 5.**
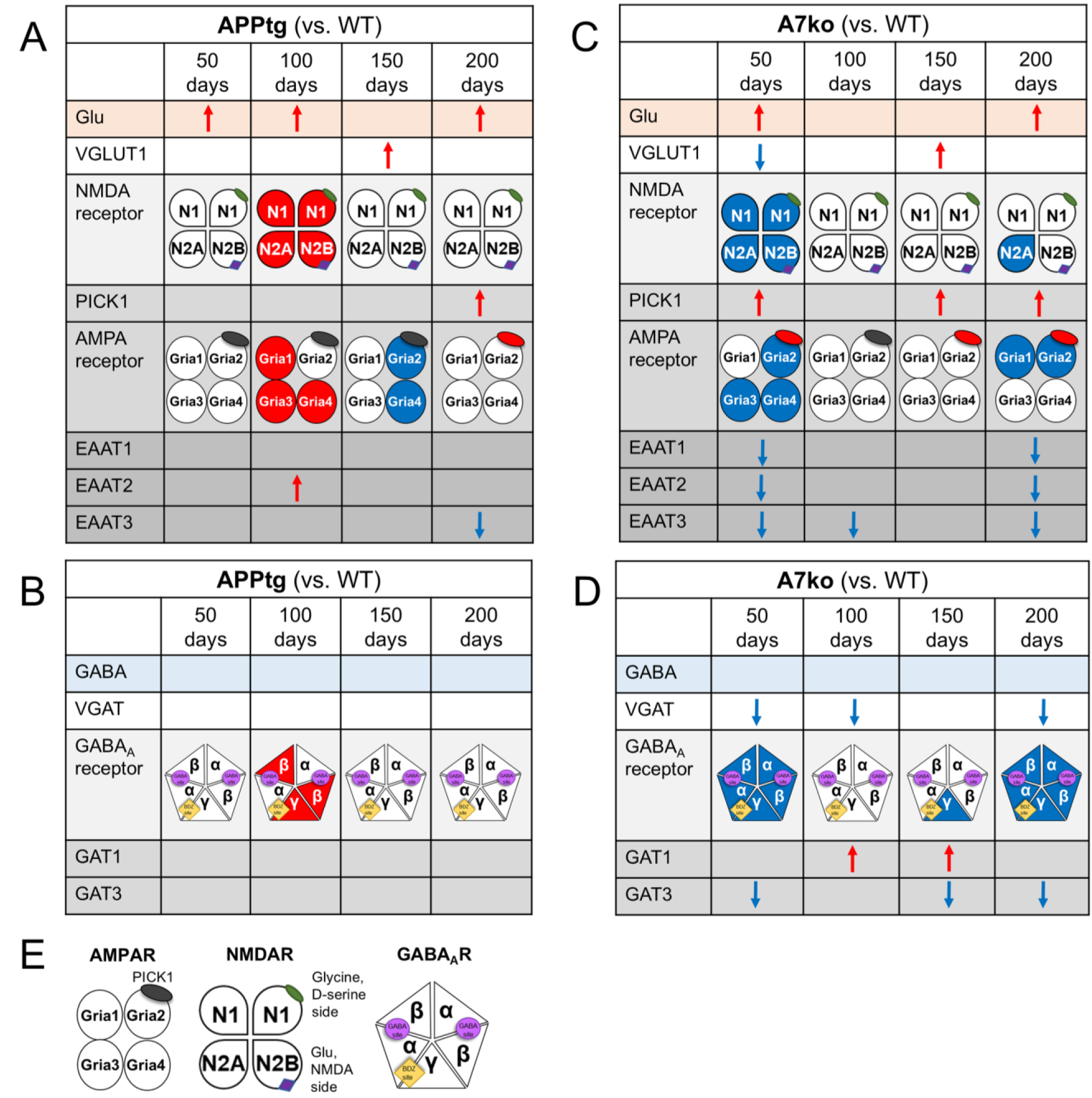
Summary of metabolomics and proteomics of the turnover of Glu and GABA in pre- and post-synaptic terminals in (A, B) APPtg vs. WT, (C, D) A7ko vs. WT in 50, 100, 150, and 200 days of age. (E) AMPAR with PICK1 binding side on GRIA2-AMPAR, NMDAR and GABA_A_R subunits composition with agonists’ binding sides. Arrows indicate increase (**red**) or decrease (**blue**) in comparison to experimental groups. Same color representation: increase – red, and decrease – blue was applied into receptor subunits composition. More details are explained in the text.

In APP-A7ko mice, a decrease in GOT1 and GLUD levels was observed in both 50- and 200-day-old animals compared to WT controls and APPtg mice (Fig. 4A and B). Similarly, in APPtg mice, the level of GLUL remained unchanged (Fig. 4D), indicating inhibition of Glu metabolism at the astrocytic level, which favors neuronal reuptake. This finding may be supported by a reduction in GLS activity during the early stage of the disease (100 days) (Fig. 4E), along with elevated Glu levels (Fig. 4C). The absence of ABCA7 in APPtg mice also impacts the GABAergic system. In the early disease stage (50 days), only an increase in GAD2 was observed (Fig. 4H). As the disease progresses, an increase in GABA levels rise at 150 days (Fig. 4I), marking a turning point in disease progression. At the same time, GAD1 levels decrease, likely due to the high GABA levels (Fig. 4G). In the late stage, reductions in GAD1, GAD2, and ABAT may result from decreased GABAergic activity and excessive Glu excitation (Fig. 4G, H, J).

A decrease in the Glu and GABA cycle of synthesis and metabolism was observed in A7ko mice. Notably, at 50 and 200 days of age, these animals showed an increase in Glu without changes in Gln and GABA (Fig. 4C, F, I). Changes in enzyme levels related to degradation were observed throughout their lifetime (from 50 to 200 days) (Fig. 4A, B, D, E, G, H). However, a reduction in metabolism at the astrocytic level was seen from early life to late stages (decreases in GOT1, GOT2, GLUL), while in neurons (decreases in GLS, GAD1, GAD2) and in astrocytes at late age (200 days) (Fig. 4A, B, D, E, G, H).

### 3.4 Dysregulation of Glu and GABA turnover in *pre- and postsynaptic terminals* is induced by Aβ deposition and lack of ABCA7

Deposing Aβ plaques directly impairs postsynaptic terminals, leading to prolonged Glu transmission due to impaired reuptake and overstimulation of receptors.

In APPtg mice, an increase in Glu levels was observed from the initial deposition of Aβ plaques in the brain, followed by a higher level of AMPAR and NMDAR subunits in 100-day-old animals (Fig. S8, S9). Also, in 100-day-old animals, an increased level of the EAAT2 transporter occurs, which may be caused by the higher level of Glu in the synaptic cleft (Fig. 5A, Fig. S5B). EAAT2 is localized mainly in astrocytes and is responsible for removing 90% of extracellular Glu. The observed changes suggest that the initial deposition of Aβ plaques initiates an increase in Glu levels, which is followed by higher activation of postsynaptic receptors and an increase in the levels of transporters responsible for removing excess Glu from the synaptic cleft. In the late stage, a decrease in the EAAT3 transporter was observed (Fig. S5C). EAAT3 is explicitly localized in neurons, with expression levels approximately 100-fold lower than those of EAAT1 and EAAT2. Additionally, EAAT3 is recruited into the membrane in the event of a Glu spill or prolonged presence of Glu in the synaptic cleft. The decrease in EAAT3 in the late stage of the disease may result from manageable levels of Glu for other EAATs in the synaptic cleft, leading to internalization of these transporters and a lack of docking at the membrane (Fig. S5C). Notably, EAAT3 is particularly concentrated in the hippocampus, targeting somatic and dendritic locations on neurons ^71^. A rare presynaptic location is restricted to GABAergic neurons, where EAAT3 provides Glu as a precursor for GABA synthesis. Evidence shows that reducing EAAT3 leads to a loss of hippocampal GABA levels and hippocampal hyperactivation ^72^. In APPtg animals, the GABAergic system was not affected. However, an increase in the β and γ subunits of the GABA_A_ receptor was observed in 100-day-old mice (Fig. 5B, Fig. S10B, C), which may be a result of the high activity of the Glu system at that same time point and the demand for GABA’s inhibitory action (Fig. 6A).

**Fig. 6.**
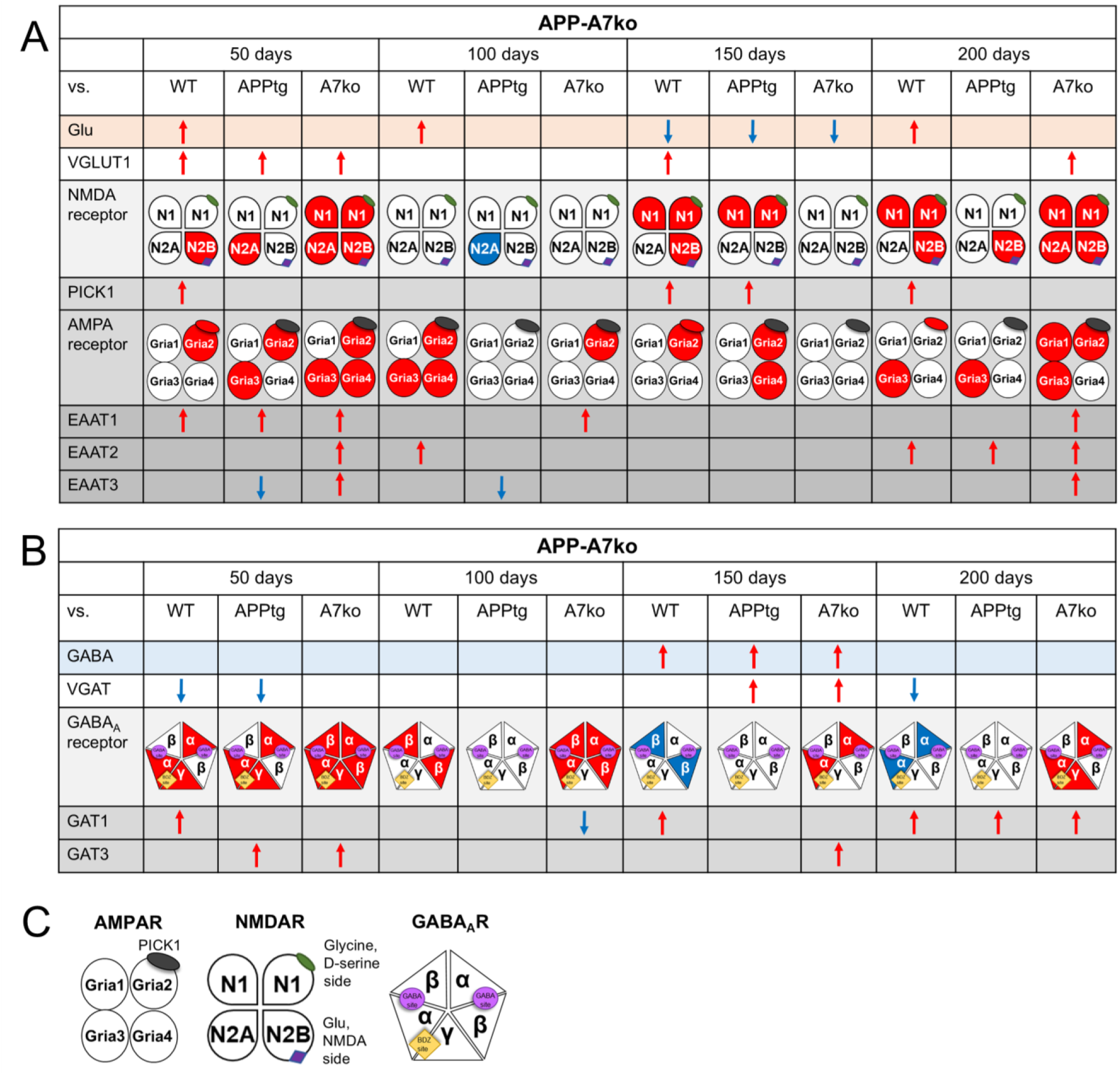
Summary of metabolomics and proteomics of the turnover of Glu and GABA in pre- and post-synaptic terminals in (A, B) APP-A7ko vs. WT, APPtg and A7ko mice in 50, 100, 150, and 200 days of age. (C) AMPAR with PICK1 binding side on GRIA2-AMPAR, NMDAR and GABA_A_R subunits composition with agonists’ binding sides. Arrows indicate increase (**red**) or decrease (**blue**) in comparison to experimental groups. Same color representation: increase – red and decrease – blue was applied into receptor subunits composition. More details are explained in the text.

A knockout of ABCA7 completely reduces the activity of Glu and GABA systems, causing a low activation state in 50- and 200-day-old animals, with less severe changes seen in 100- and 150-day-old mice (Fig. 6C and D). In these mice, the increase in Glu levels at 50 days was linked to a decrease in NMDAR subunits (Fig. S9). Additionally, the absence of ABCA7 simultaneously lowers AMPA receptor subunit 2 (GRIA2) (Fig. S8B) and raises PICK1 (Fig. S8E) (shown by arrows indicating the decrease of GRIA2 with the increase of PICK1 in the A7ko group – blue bars). PICK1 (protein interacting with kinase C1) is involved directly in AMPAR internalization (68), forming a protein-protein complex with the receptor’s second subunit (GRIA2). PICK1 directly causes the reduction of surface GRIA2 AMPAR, signaling the internalization of AMPA receptors, which leads to a loss of synaptic communication and strength. NMDAR and AMPAR are crucial in facilitating long-term potentiation (LTP), a key example of synaptic plasticity that supports neuronal development and maturation (69). It is well established that Glu and GABA systems are vital for CNS development, and the observed low activation at 50 days may delay this process. The decreased synaptic transmission is supported by low levels of all EAAT transporters, indicating a lack of Glu in the synaptic cleft and possibly suggesting Glu is stored in synaptic vesicles inside glutamatergic neurons. A similar low activity state in the GABA system and impaired GABA release are also observed, which can simply result from a lack of excitatory activity. In 50-day-old mice, decreased levels of the vesicular transporter (vGAT) (Fig. S6C) and the reuptake transporter GAT3 (Fig. S6B) were found, along with reduced expression of GABA_A_ receptor subunits (Fig. S10).

Both systems are less affected in 100- and 150-day-old animals compared to the early and late time points (Fig. 6C and D). In these animals, no changes in EAAT transporters 1 and 2 were observed, with a decrease in EAAT3 (Fig. S5A, B, C) and receptors (Fig. S8 and S9), along with an increase in the GABA reuptake transporter GAT1 (Fig. S6A). The less severe changes may result from the maturation of the animals. However, the impact of ABCA7 on the Glu and GABA systems is profound, leading to a decrease in the activity of all excitatory transporters (Fig. S5A, B, C), a reduction in NMDAR subunit levels (Fig. S9), and PICK1-related AMPAR internalization (Fig. S8B and E). Additionally, there is a decrease in the activity of the GABAergic system, evidenced by lower levels of VGAT1 and GAT3 transporters (Fig. S6B and C) and a reduced expression of GABA_A_ receptor subunits (Fig. S10).

In 50-day-old APPtg mice, the lack of ABCA7 increases the activity of VGLUT1 (Fig. S5D) and the astrocytic EAAT1 transporter (Fig. S5A) in response to high Glu levels (Fig. 6A). With the progression of the disease to 150 days, a lower level of Glu is followed by an increase in the N1 and N2B subunits of NMDAR (Fig. S9A and C) as well as GRIA2 and GRIA4 subunits of AMPAR (Fig. S8B and D) (Fig. 6A). At the later time point of 200 days, characterized by high levels of Aβ in the brain, an increase level of Glu is concomitant with an increase in EAAT2 levels (Fig. S5B), the leading astrocytic transporter, as well as NMDAR N1 and N2B subunits (Fig. S9A and C), suggesting a high concentration of Glu in the synaptic cleft (Fig. 6A). In APPtg mice lacking ABCA7, an increase in GABA_A_R subunits (Fig. S10A and C) occurs alongside an increase in GATs (Fig. S6A and B) at early stages, possibly as a compensatory mechanism for high Glu system activation (Fig. 6B). Interestingly, at 150 days, the GABA system counteracts changes observed in the glutamatergic system, with an increase in GABA levels and a decrease in Glu levels, suggesting a predominance of inhibitory neurotransmission over excitation as a potential rescue mechanism due to progressive Aβ deposition (Fig. 6A and B). At the later time point, an overdrive of excitatory neurotransmission diminishes GABAergic inhibition, with only an increase in GAT1 observed (Fig. S6A), which may serve as a compensatory mechanism in response to the lack of inhibitory activity (Fig. 6B).

### 3.5 Aβ deposition and the lack of functional ABCA7 primarily affect the levels of LPC and LPE in the brain of aged animals

Alterations in brain lipid metabolism and accumulation are a consistent hallmark of AD and are increasingly regarded as part of its molecular signature. Here, we performed heat map and PCA analyses demonstrate that changes in the lipidome are observed in aged mice, where the progressive deposition of Aβ plaques (APPtg and APP-A7ko) leads to an increase in lipid levels (Fig. 7C and D). Separate PCA analyses were performed to evaluate which lipid classes are most affected by AD pathology and lack of ABCA7 (data not shown). Based on the PCA results, the LPC and LPE lipid classes were identified as the most affected, which was further confirmed by Volcano plot analysis (Fig. 8A, B, C).

**Fig. 7.**
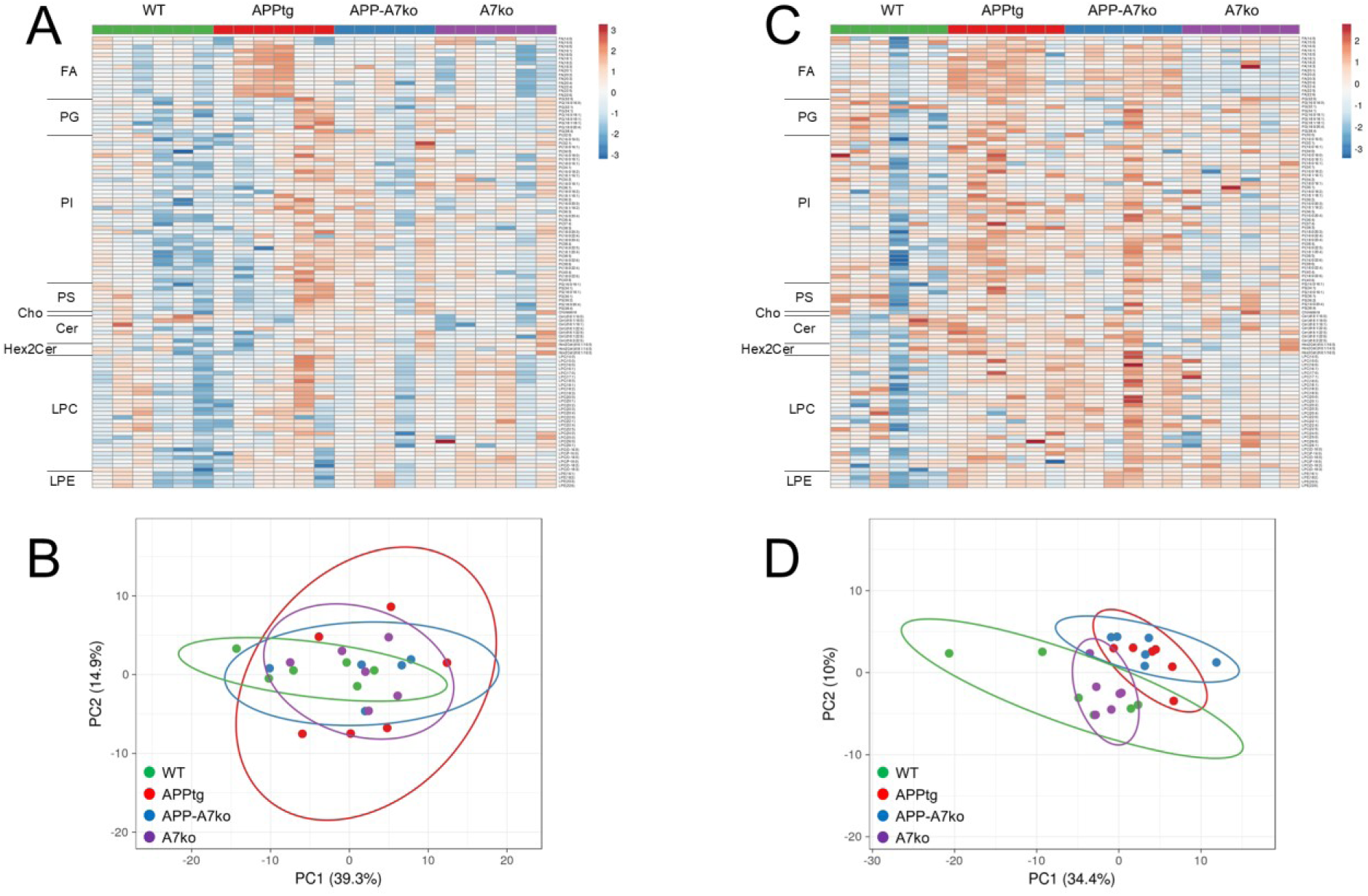
Deposition of Aβ increases lipid accumulation in the brains of aged animals. Heat map analysis of lipid classes: FA, PG, PI, PS, Cholesterol, Cer, Hex2Cer, LPC, and LPE in (A) 50-day-old and (C) 200-day-old WT control mice, APPtg mice, APPtg mice with ABCA7 transporter deficiency (APP-A7ko), and animals lacking ABCA7 (A7ko). Principal component analysis (PCA) plots show the percentage of variability between samples due to genotype effects in (B) 50-day-old and (D) 200-day-old mice. All analyzed lipids, in the order presented in the heat maps, are listed in the Supplementary Materials (Fig. S14).

**Fig. 8.**
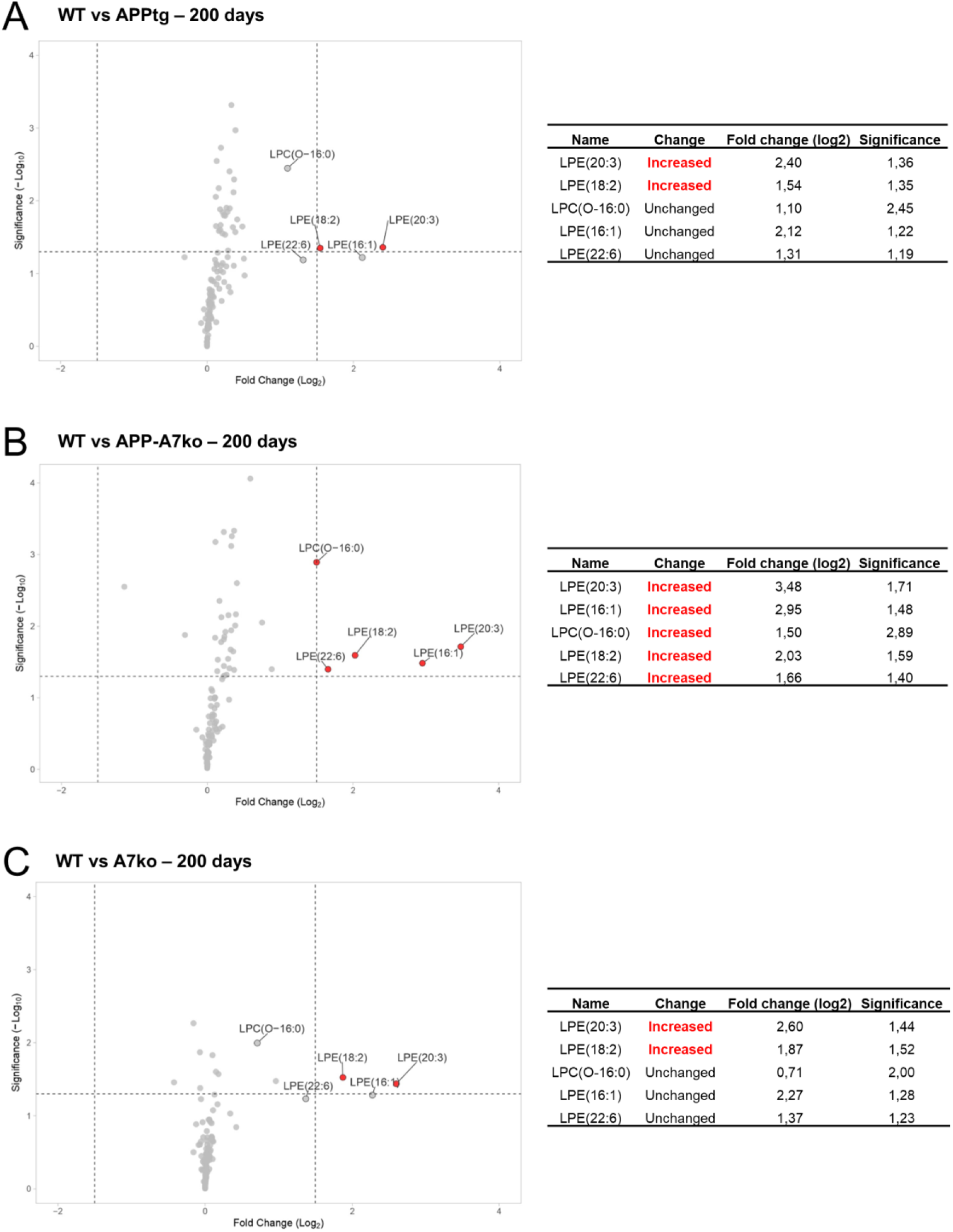
Aβ deposition and the lack of ABCA7 primarily affects LPC and LPE levels in the brains of aged animals. The volcano plot illustrates changes in lipid levels between (A) WT vs. APPtg, (B) WT vs. APP-A7ko, and (C) WT vs. A7ko at the age of 200 days. The horizontal dotted line represents a significance level of p = 0.05 (–log_10_ = 1.3), and the vertical dotted lines indicate a fold change of ±1.5 (log_2_).

Most significant changes were observed in the levels of LPC (Fig. S15A and B) and LPE (Fig. S15C and D), with increased accumulation depending on progressive Aβ deposition and the lack of ABCA7 (200 days of age). Based on the volcano plot data, both factors: an increase in Aβ levels along with the knockout of ABCA7 lead to higher accumulations of four LPE species: LPE(20:3), LPE(16:1), LPE(18:2), LPE(22:6), and one LPC species (LPC(O-16:0)) (Fig. 8B). When these factors occurred separately, only two LPE species, LPE(20:3) and LPE(18:2), remained elevated (Fig. 8A and C). These results highlight LPE(16:1), LPE(22:6), and LPC(O-16:0) as lipids that are increased explicitly in APP-A7ko mice. Moreover, LPE(20:3) is a lipid that is upregulated across all experimental groups, with the highest level observed in APP-A7ko mice (Fig. 8A, B, C).

### 3.6 Virtual lipid landscape analysis using MSI-ATLAS

The recently described MSI-VISUAL ^42^ and MSI-ATLAS ^44^ frameworks provide computational and machine learning tools to visualize and determine the region-specific lipid composition based on Mass Spectrometry Imaging data of the brain.

Using brain lipidomics data, followed by a highly detailed brain segmentation model, a Computational Brain Lipid Atlas (CBLA) and Virtual Landscape Visualizations (VLV) of the lipid composition across different brain regions and their anatomical connections were generated. Through VLV, we present regions with the highest accumulation of lipids based on their mass-to-charge ratio (m/z). The resulting color maps represent lipid accumulation levels corresponding to signal intensity, ranging from 0 (blue) through green, yellow, and brown to white (maximum 255, indicating high accumulation).

Here we presented only the localization of LPE(22:6) (502.30 m/z) in the AD mouse brain with or without ABCA7 transporters, because this lipid is specifically increased in white matter regions of APP-A7ko animals (Fig. 9). In human post mortem tissue coming from AD individuals with *PSEN1* mutation LPE(22:6) was also observed in the plaques, however despite the enrichment this species was not significantly increased in Aβ plaques on a group level (2 out of 5 patients) ^73^.

**Fig 9.**
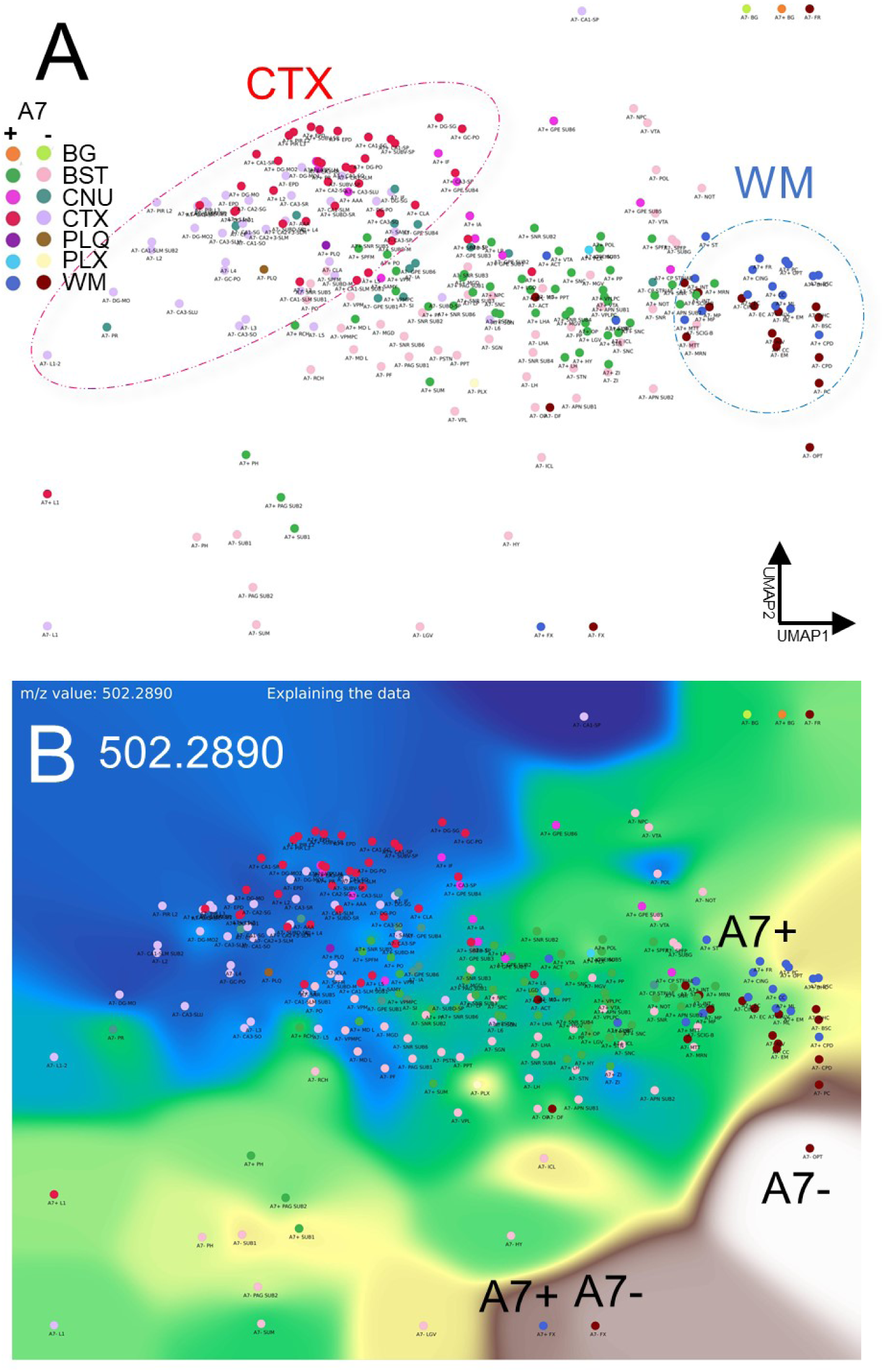
Virtual Landscape Visualization (VLV) of a differentially expressed m/z in AD mice (A7+) versus AD-A7ko mice (A7-). (A) shown is the UMAP 2D dimensionality reduction graph of 123 brain regions highlighting the cortex (CTX) and white matter (WM) region clusters. A7ko mice show a shift in their lipid profile. (B) Virtual landscape visualization the m/z 502.2890 showing increased levels in AD-A7ko mice (A7-) in white matter regions. No expression is detected in cortex regions (blue VLV). The color maps represent signal intensity, ranging from 0 (blue) through green, yellow, brown, to white (maximum 255) (Matplotlib, color map: terrain). Brain area nomenclature is based on the Allen Mouse Brain Atlas (https://mouse.brain-map.org/).

### 3.7 Locomotor activity

In mice, spontaneous locomotor activity reflects circadian rhythm and serves as a marker of normal neurological and behavioural function. It also indicates the drug’s potential to restore healthy rhythms. Sleep and circadian disturbances are early features of AD, often appearing before cognitive symptoms. Fragmented sleep, reduced deep sleep, and disrupted day–night activity patterns have been observed in preclinical stages, making them promising early, biomarkers of disease progression ^74^. Based on this, we applied this behavioral test to investigate whether there is a correlation between disease progression, ABCA7 loss of function, and alterations in mice locomotor activity patterns and circadian rhythms.

In animals with Aβ deposition, both with and without ABCA7, a similar pattern of locomotor activity was observed in the early stages of the disease (APPtg and APP-A7ko, Fig. 10A and B, purple line). As the disease progressed, APPtg mice exhibited increased and sustained activity (Fig. 10A, bold red line on days 2 and 3). However, in AD animals lacking ABCA7, inhibition of locomotor activity was noted (Fig. 10B, bold green line). In control animals, the lack of ABCA7 inhibited locomotor activity from the early time point (50 days) through the later stages of the disease (Fig. 10C, purple line for 50 days and bold blue line for 200 days).

**Fig. 10.**
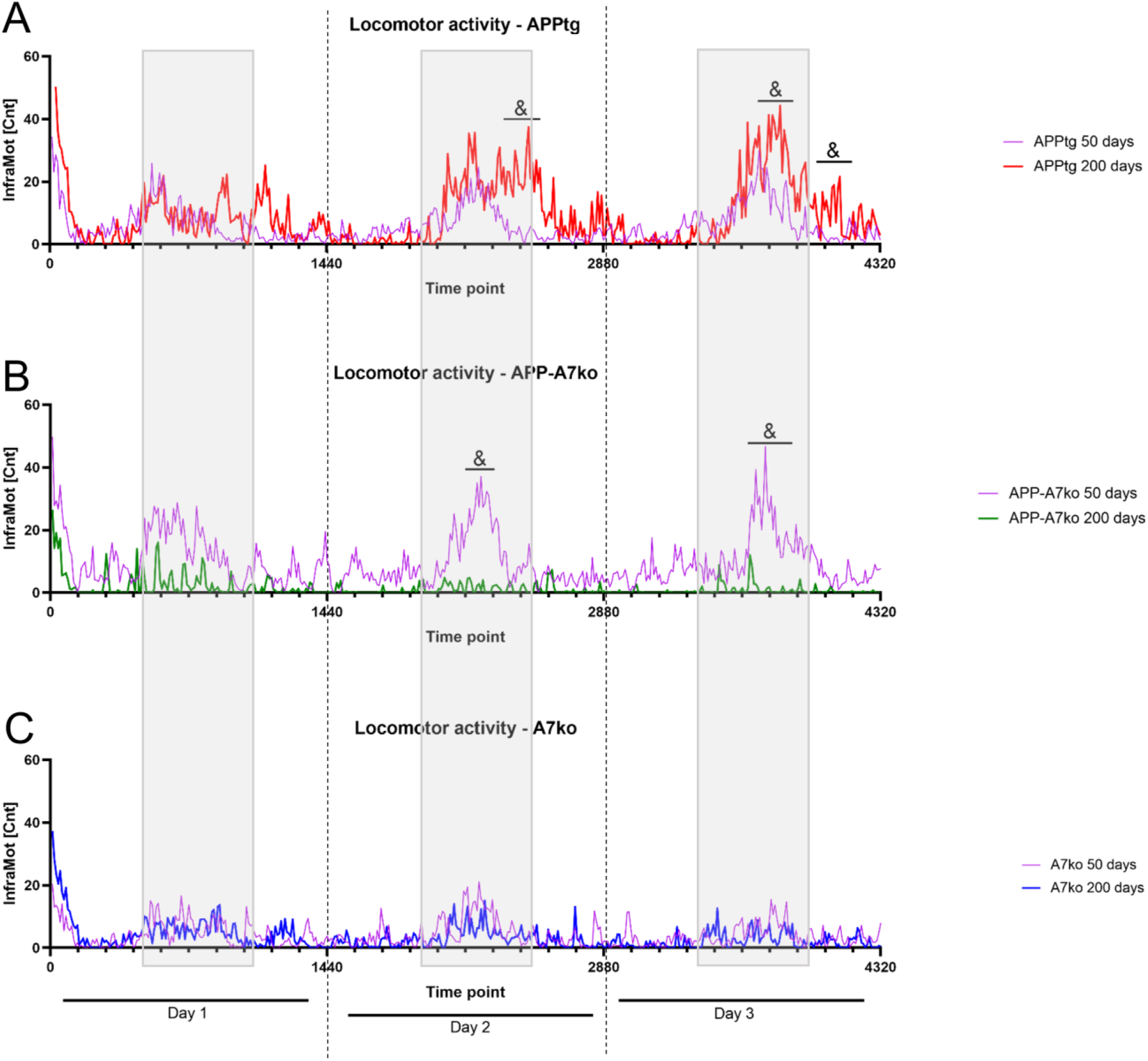
Lack of ABCA7 decreases locomotor activity. Differences in spontaneous locomotor activity between mice at 50 days old (purple line) and 200 days old (bold line) are shown for (A) APPtg (red), (B) APP-A7ko (green), and (C) A7ko (blue). Activity was measured every 10 min over 72 h, with each day consisting of 1440 min. The locomotor activity pattern for each group is presented as mean values. Gray rectangular represent the dark (active) phase of the circadian cycle. Statistical analyses were performed using a two-tailed unpaired Student’s t-test after a normality check; p < 0.05 was considered significant for comparisons between 50 days and 200 days.

## 4 Discussion

In this study, we present a comprehensive analysis of the Glu-Gln-GABA system in Alzheimer’s disease (APPtg) mice, both with and without ABCA7, throughout the progression of the disease.

We analyzed proteomic data of all astrocytic and neuronal markers, including degrading enzymes, transporters and receptors, as well as the metabolites Glu, Gln, and GABA. This analysis reveals the changes in neurotransmitters turnover and the mutual communication between astrocytes and neurons, highlighting the crucial cooperation required to balance excitatory and inhibitory neurotransmission. An imbalance between these systems can have lasting consequences for the brain’s neuronal networks. Furthermore, by analyzing specific lipid classes, we identified two lipids classes (LPC and LPE) that were significantly upregulated in APPtg mice lacking ABCA7.

The primary aim of the studies presented here was to describe how ABCA7 transporters affect the Glu-GABA imbalance caused by AD pathology. To investigate this, we used the well-established APP/PS1 mouse model of AD, characterized by the progressive deposition of Aβ plaques in the brain, starting in the cortex and extending to the hippocampus, striatum, thalamus, and brainstem ^45^. In these animals, a linear increase in Aβ load is observed beginning at 3 months of age. Using this model, we detected the first plaques in the cortex of 50 days old mice (approximately 1.5 months old). In our findings, the increase in brain Aβ load corresponds with a decrease in CSF levels starting at 6 months of age, a crucial time point in disease progression ^75^. These results align with our analyses, which show an increase in APP at 150 days of age (5 months old) compared to earlier stages of the disease (up to 100 days), with APP levels remaining stable until late-stage (200 days of age). Additionally, we observed elevated levels of APP in other CNS compartments, such as ISF and CSF. In this context, the CSF reflects changes in the brain, where the progressive increase in APP, coupled with the loss of BBB integrity, contributes to APP accumulation in the CSF. Furthermore, APP was also detected in the ISF, which represents the extracellular space. The presence of APP, a membrane protein, suggests possible disruptions in cellular membranes, highlighting a central pathological mechanism of AD induced by Aβ peptides ^76,77^.

### 4.1 Aβ plaque deposition facilitates Glu-related toxicity

The critical role of Glu-GABA system imbalance in the pathogenesis of AD is well established and represents a potential target for treatment. In the brains of AD patients, decreased level of Glu are associated with neuronal loss ^78,79^. Furthermore, meta-analyses indicate that AD pathology leads to reductions in Glu levels, Glu reuptake, and specific subunits of NMDAR, N2B, and AMPAR, Gria2/3 ^80^.

In APPtg mice, an increase in Glu was correlated with reduced activity of GLUD1. GLUD1 is enriched in regions with high glutamatergic neurotransmission and is an enzyme that reversibly converts Glu into αKG ^81^. These early changes also influence the GABAergic system; the higher activity of GAD2 (GAD65) may indicate that Glu is used as a precursor for GABA to prevent Glu-related overstimulation. The GABA system is significantly affected in AD ^82^. In GABAergic neurons, GABA is synthesized by two GAD isoforms: GAD1 (GAD67) and GAD2 (GAD65). Two distinct pools of GABA have been described in the brain: a metabolic pool and a neurotransmitter pool, with the localization of GAD isoforms correlating with these pools. GAD65, which supplies the neurotransmitter (vesicular) pool, is preferentially found in neuronal terminals and is considered the primary source of GABA in response to synaptic demand ^83,84^. In contrast, GAD67 is located in the cytoplasm of neurons and produces GABA for the metabolic pool. Importantly, this differentiation is not definite, as both GAD isoforms can contribute to both pools ^85^.

Moreover, in our studies AD progression is associated with increased activity of EAAT2 transporters, likely due to elevated concentrations of Glu in the synaptic cleft, leading to system adaptation. EAAT2 can retract 90% of extra-synaptic Glu ^86^, thus playing an essential role in maintaining Glu homeostasis and preventing Glu-related excitotoxicity. EAAT2 is expressed at high levels in astrocytes and at lower levels in neurons ^87^. Numerous human and animal studies have indicated impairments in EAAT2 function in the pathogenesis of AD. In AD patients, the expression of EAAT2 in the hippocampus and medial frontal lobe is decreased ^88–90^. In murine models of AD, loss of EAAT2 accelerates the development of cognitive impairment ^91,92^. Restoration of EAAT2 function using a novel EAAT2 translational activator has been shown to improve cognitive function ^93^. Other studies have demonstrated a direct correlation between increased Aβ load and Glu neurotoxicity via impaired EAAT2 function. In primary neuronal-astrocyte co-cultures, Aβ and Glu have been found to decrease both cell viability and EAAT2 levels. Restoring EAAT2 function with Sulbactam, as evidenced in a rat model of brain ischemia ^94^, protects cells against concomitant Aβ and Glu toxicity through upregulation of EAAT2 expression ^95^. However, some studies indicate a lack of changes in EAAT2 levels in the brains of individuals with advanced AD. In these cases, mRNA levels of EAAT2 were preserved, although there was a tendency for EAAT2 protein levels to decrease in the frontal cortex. This may suggest adaptations to neuronal dysfunction rather than alterations in EAAT2 expression ^96^. There is a substantial amount of data indicating that EAAT2 levels are regulated by various factors, including synapse development and maturation ^97^ and astrocyte plasticity (astrocyte coverage), which depend on neuronal activity and synaptic release ^98,99^.

Overstimulation of postsynaptic NMDA and AMPA receptors is a crucial aspect of prolonged Glu accumulation in the synaptic cleft. In our studies, increases in all NMDAR subunits, as well as subunits 1, 3, and 4 of AMPAR, were observed beginning from 100 days of age. The presence of these receptor subunits was also detected in CSF, suggesting either degradation of postsynaptic terminals or an increase in turnover and docking of the receptors. Our results are in line with studies showing presence of NMDAR – GluN2A, GluN2B, GluN1, and GluN3A subunits level in CSF coming from AD patients, where in AD CSF the level of GluN2B was significantly decreased. Importantly, in those studies CSF samples were obtained from individuals seeking medical evaluation for cognitive impairment ^100^. It has been shown that Aβ plaques co-localize with the GluN1 and GluN2B subunits of NMDA receptors on neuronal soma and neurites ^101^. NR2A subunits are predominantly located at the synapse, and their activation is associated with the expression of pro-survival genes. Conversely, NR2B subunits are primarily located extrasynaptically and have been linked to cell death and neurodegeneration ^102,103^. Therefore, activating NR2A and/or inactivating NR2B-containing NMDARs has been proposed as a therapeutic strategy for treating AD. Research has demonstrated that NR2A-containing NMDARs decrease neuronal death and promote cell viability following Aβ treatment, while NR2B-containing NMDARs exhibit higher levels of cytotoxicity and lower levels of neuronal survival ^104^.

### 4.2 ABCA7 deficit promotes AD pathogenesis due to Glu-GABA imbalance

In the context of AD, ABCA7 is implicated in lipid transport as well as the clearance of toxic Aβ. However, little is known about how a lack of ABCA7 affects neurotransmission. Impaired function of ABCA7 may contribute to AD pathology through two primary mechanisms: First, ABCA7 facilitates the processing of APP and increases Aβ deposition ^31,35^. Second, ABCA7 dysfunction impairs the ability of microglia to phagocytose Aβ ^36^. Our results indicate that the absence of ABCA7 in APPtg mice increases Glu-related toxicity, likely due to astrogliosis rather than enhanced Aβ production. In APP-A7ko mice, there were no differences in AD-related protein levels across all examined compartments (whole brain, CSF and ISF) when compared to APPtg mice. However, the level of GFAP was found to be elevated starting from 150 days of age compared to both control and APPtg mice. Furthermore, our previous studies showed that the knockout of ABCA7 transporters in APP/PS1 mice resulted in an increase level of Aβ42, which correlates with higher activation of microglia and astrocytes, as evidenced by an increased number of IBA1 and GFAP-positive cells, respectively ^46^.

In APPtg mice lacking ABCA7, the imbalance between Glu and GABA is more profound compared to APPtg mice. In glutamatergic system, an increase in neuronal EAAT1 and vGLUT1 was observed, suggesting a higher turnover of Glu that bypasses astrocytic metabolism. Additionally, elevated levels of NR2A and NR2B subunits of NMDA receptors indicate an increase in neurotransmission and a possible co-localization of NR2B with Aβ, which may exacerbate neurotoxicity ^102,103^. Simultaneously, activation of the GABAergic system may counterbalance the diminished excitatory glutamatergic signaling. The damaging effects of ABCA7 deficiency become particularly profound from 150 days of age, a critical time point in disease progression. In contrast to APPtg mice, where we observed some adaptation in response to the disease, the Glu-GABA imbalance persisted without significant progression. At the peak of the disease, the inhibitory neurotransmission appears to overtake excitation, characterized by increased GABA levels and decreased Glu levels. This suggests that the system may be attempting to compensate for the concomitant overactivation of Glu. A significant aspect that supports our findings is the increased expression of NR1 and NR2B subunits of NMDA receptors. These subunits contain binding sites for Glu and the co-agonist glycine ^105,106^. While the role of the NR2B subunit in relation to Aβ plaques has been previously discussed, it is also important to note that the glycine-binding site is located on the NR1 subunit. The binding of both glycine and Glu is necessary for the removal of the Mg²⁺ block and subsequent stimulation of NMDA receptors ^107^.

In the late stage of AD, the entire neurotransmission system becomes overdriven by Glu. The impairment of Glu metabolism in astrocytes, combined with increased activity of EAAT2, may enhance the reuptake of Glu into astrocytes, leading to increased astrogliosis observable in all compartments. The elevated levels of GFAP in the ISF and CSF suggest the extreme activation or destruction of astrocytes in the brain leading to GFAP leakage into the periphery. Prolonged activation of Glu neurotransmission results in impairment of the inhibitory GABAergic system. The observed increase in GAT1, along with decreased vGAT and GAD1/GAD2, indicates an adaptation or compensation for the lack of GABA neurotransmission. A decrease in the α-subunit of the GABA_A_ receptor further supports this. Research by Schwab et al. shows that deficiencies in GAD65/67 contribute to the loss of GABAergic inhibitory activity ^84^. Our findings align with meta-analysis data reporting decreases in GABA, GAD65/67, GABA transporters, and GABA_A_ receptors in the brains of individuals with AD ^108^. Additionally, direct evidence demonstrates that forming Aβ plaques can reduce the number and activity of GABA interneurons in the hippocampus, leading to cognitive impairments ^109^. Aβ also downregulates the surface levels of the GABA_A_ receptor, thereby weakening GABA inhibition ^110^.

### 4.3 Lack of ABCA7 diminished the function of the Glu and GABA systems

To date, there is limited data on how the absence of ABCA7 affects the Glu-GABA systems. In African Americans, the ABCA7 variant rs115550680 is the strongest genetic locus associated with late-onset AD, and research has shown that healthy carriers of this variant exhibit impaired cortical connectivity ^111^. Additionally, the presence of the rs3764650G allele in ABCA7 has been linked to the development of neuritic plaque pathology ^112^ and to ABCA7 deficiency ^31^, which is associated with lower memory scores ^113^. In animal studies, homozygous Abca7 knockout mice (Abca7−/−) demonstrated sex-specific deficits in certain cognitive functions; male mice showed impairments in novel object recognition memory, while female mice exhibited deficits in spatial reference memory. Furthermore, acute treatment with the NMDAR antagonist MK-801 in ABCA7-deficient female mice produced a locomotor-stimulating effect, which may indicate increased sensitivity or expression levels of NMDARs ^114^.

In the study presented here, the absence of ABCA7 led to a complete decline in the activity of the Glu and GABA systems. Throughout early life, there was a noted decrease in the expression of transporters and receptors, continuing into late age, where all examined components of the Glu-GABA system were diminished. Interestingly, despite these decreases, Glu levels in the whole brain remained high both in the early and late stages. It is well-established that glutamatergic neurotransmission plays a critical role in CNS maturation from early life, primarily through NMDA-AMPA-dependent synaptic plasticity ^115^. For synaptic maturation, the insertion of both AMPA and NMDA receptors is crucial ^116^, along with a developmental switch in NMDAR subunit composition from NR2B to NR2A ^117^. This switch is evolutionarily conserved and occurs in various brain regions, including the cortex, hippocampus, amygdala, and cerebellum. In our control mice with a knockout of ABCA7 transporters, we observed a decrease in the levels of NMDA and AMPA receptors, alongside an increase in PICK1 protein levels, which is involved in the internalization of AMPA receptors ^118^. The downregulation of surface AMPA receptors reduces synaptic communication, leading to cognitive impairments in neurodegenerative diseases such as AD ^119^. Notably, it has been shown that knockout of PICK1 or blocking the PDZ domain of PICK1 with BIO922 can inhibit the interaction between GluA2 and PICK1, preventing the loss of AMPAR-mediated synaptic transmission and the reduction of AMPAR surface levels caused by Aβ ^119^.

### 4.4 Lack of ABCA7 in APPtg and WT mice decreases locomotor activity

In our studies, the knockout of ABCA7 in healthy and APPtg mice resulted in a decrease in spontaneous locomotor activity in the dark (active) phase. However, in APP-A7ko mice lower activity was associated with glutamatergic system overactivation in late stage of the disease (200 days), while in A7ko group with diminished Glu-GABA system that was reported in 50 and 200 days. In AD individuals that disturbance in circadian rhythm is one of the first features of AD in patients ^120^, thus using this test in rodents may be a good indicator of disease progression and evaluation of possible new treatments efficacy.

Till now, there is not many studies investigating Glu-GABA system alteration and changes in locomotor activity in rodents. Research indicates that restoring GABA activity through GABA agonists may help mitigate the progression of motor symptoms in Parkinson’s disease (PD) ^121^. In contrast to the GABA system, increasing the expression of the Glu transporter EAAT2 with ceftriaxone improved Glu reuptake, which in turn enhanced locomotor function in a PD model ^122^. Moreover, elevated EAAT2 activity has been shown to attenuate morphine-conditioned place preference, morphine-induced hypothermia, and locomotor sensitization ^123^. Riluzole, a drug proven to restore EAAT2 expression in the nucleus accumbens, has been effective in preventing cocaine reinstatement ^124^ and reducing methamphetamine-induced locomotor sensitization ^125^. The role of glutamatergic neurotransmission in locomotor activity is further supported by findings that NMDA receptor antagonists, such as MK-801 and ketamine, can induce psychostimulant effects ^126,127^. In our APPtg mice lacking ABCA7, we noted an increase in NMDA receptor expression alongside higher levels of EAAT2. However, it is challenging to draw definitive conclusions about how changes in the Glu-GABA system led to decreased locomotor activity. We hypothesize that this decrease may be attributed to overexcitation of the glutamatergic system, resulting in heightened Glu-related neurotoxicity.

### 4.5 Lipidomics disease signatures

Our lipidomics analysis revealed that AD pathology primarily affects the LPE and LPC lipid classes. Notably, the absence of ABCA7 in APPtg mice resulted in a distinct lipidomics signature associated with the disease. In these mice, we observed increased levels of LPC(O-16:0), LPE(16:1), and LPE(22:6). Additionally, LPE(20:3) was upregulated across all experimental groups, with the most significant accumulation observed in mice exhibiting both Aβ plaque deposition and a lack of ABCA7. These findings were further validated by landscape analysis of LPE(22:6), which demonstrated its specific localization in the brain’s white matter regions.

LPE and LPC are polar phospholipids generated by phospholipase A2 (PLA2) from phosphatidylethanolamine (PE) and phosphatidylcholine (PC), respectively ^128^. Under pathological conditions, reactive oxygen species (ROS) can metabolize phospholipids into their lyso-forms ^129^, which, alongside Aβ plaques, present important features of neurotoxicity. It is well established that in AD, antioxidant defense mechanisms are impaired, leading to increased ROS production due to Aβ aggregation ^130^. However, it remains unclear whether alterations in phospholipid metabolism are a consequence of Aβ deposition or if changes in lipid metabolism trigger abnormal aggregation. Studies have shown that PLA2 activity is specifically increased in astrocytes in the cortex of AD patients and in animal models of AD ^131–133^, indicating that neurodegeneration in this area may be linked to lipid metabolism ^134^. In our studies, the concomitant increase in Aβ plaque deposition and Glu toxicity may also affect the accumulation of LPC and LPE through PLA2 overactivation. The connection between PLA2 activation and Glu toxicity can be attributed to the elevation of calcium concentrations resulting from NMDA receptor activation ^135^. Additionally, LPE has been shown to induce calcium influx in PC-12 neuronal cells and SH-SY5Y neuroblastoma cells ^136^.

In our analysis, LPE(20:3) was upregulated in all experimental groups, with the highest levels observed in the group with AD pathology and a lack of ABCA7. In contrast, LPE(22:6) and LPE(16:1) were specifically elevated only in the APP-A7ko animals.

ABCA7 transporters play a crucial role in the clearance of Aβ ^34^, as well as in the release of cellular cholesterol and phospholipids ^137^. There is strong evidence linking cholesterol to Aβ production; exposure of neuronal cells to cholesterol transport-inhibiting agents has been shown to increase γ-secretase activity, leading to elevated production of Aβ40 and Aβ42 ^138^. These findings highlight the involvement of ABCA7 transporters in the development of AD and their significance in healthy individuals, while also suggesting that LPE(22:6), LPE(16:1), and LPE(20:3) may serve as indicators of disease progression related to Aβ levels. Our data align with other studies that have reported increased levels of LPE(22:6) in cases of vascular dementia ^139^. Additionally, in cognitively impaired ischemic rats, both LPE(22:6) and LPE(20:3) were found to be elevated in the hippocampus. Treatment with Linalool, an NMDA receptor inhibitor, restored lipidomics profiles and improved performance on motor and cognitive tests in these animals, highlighting the role of glutamatergic neurotransmission ^140^. Furthermore, accumulation of LPE has also been reported in the brains of AD model mice (3xTg-AD), where silencing BACE1, an enzyme involved in the amyloidogenic pathway of APP, restored the lipidomics profile ^141^. Conversely, human studies have shown lower levels of LPE(22:6) and LPE(20:3) in symptomatic AD patients compared to control and asymptomatic individuals ^142^. Animal studies have reported similarly declining levels of brain LPE(22:6) with aging and under conditions of impaired spatial cognitive abilities or memory ^143^. The importance of LPE(22:6) in AD pathogenesis is further emphasized by landscape analysis, which reveals lipid accumulation in the Aβ plaques in the optic tract (OPT), a part of white matter. Loss of white matter microstructural integrity has been reported in patients with AD, representing neural network deficits that underlie gradual cognitive decline ^144,145^. The OPT consist of the axons originating from the retinal ganglion cells (RGCs) in the retina, carry visual information from the optic chiasm to the lateral geniculate bodies as a part of the visual pathway. Post-mortem studies of AD patients have revealed RGC degeneration accompanied by AD-specific pathology, including the presence of Aβ deposits in the retina ^146^.

The involvement of LPC in AD pathology has been demonstrated, with increased LPC accumulation observed in the white matter of aged human brains exhibiting senile atrophy of the Alzheimer type ^147^. Additionally, LPC has been shown to enhance Aβ-induced neuronal apoptosis ^148^ and to promote the formation of Aβ(1-42), including its trimers and tetramers ^149^. In our studies, LPC(O-16:0) was specifically increased in AD-A7ko mice. Interestingly, this lipid has also been found in Niemann-Pick disease type C (NPC), a rare lipid storage disorder characterized by the accumulation of unesterified cholesterol, disruptions in lysosomal calcium homeostasis and signaling pathways, mitochondrial dysfunction, increased inflammation, oxidative stress, and ultimately, neuronal degeneration ^150^. In animal models of NPC, elevated levels of LPC(O-16:0) have been detected in the liver, spleen, cerebrum, cerebellum, and brainstem ^151^. Furthermore, the same study reported increased levels of alkyl-LPCs in the prefrontal cortex, hippocampus, and cerebellum of NPC1 patients ^151^. Importantly, in these patients, an increase in Aβ42 has been linked to enhanced γ-secretase activity ^152^. This finding is consistent with observations in NPC model cells, where Aβ accumulation is associated with elevated cholesterol levels in late endosomes ^153^.

### 4.6 Conclusions and future directions

The results of this study emphasize the importance of ABCA7 dysfunction in the development of AD, especially in relation to the regulation of glutamatergic and GABAergic neurotransmission. Specifically, the deposition of Aβ impaired the glutamatergic system. Notably, the absence of ABCA7 in APPtg mice led to more severe alterations characterized by an overdrive of excitatory neurotransmission and reduced inhibitory signaling.

As a lipid transporter, ABCA7 may influence the lipid content of neuronal membranes, thereby impacting neurotransmission at both presynaptic and postsynaptic levels, affecting the release of vesicles and the docking of receptors into the membrane. However, the specific mechanisms by which ABCA7 mediates these effects need further investigation.

Our lipid analysis hypothesize LPC(O-16:0), LPE(22:6), and LPE(20:3) as promising indicators of disease progression and potential treatment targets, emphasizing their specific localization in pathological contexts.

The findings also raise the question of whether existing drugs that modify glutamatergic neurotransmission in other neurological conditions could be repurposed for patients with AD. To date, the only drug primarily targeting Glu-related toxicity is memantine. This NMDAR antagonist has been used since 1986 and is considered safe and helpful for symptomatic treatment in AD ^154^. However, memantine is ineffective for all patients, with an efficacy rate of about 70% ^155^. Additionally, there are concerns regarding its diminishing effectiveness over time, as some patients may find that it becomes less effective or eventually stops working altogether.

Our study links the sole absence of ABCA7 (without any disease) to reduced glutamatergic and GABAergic neurotransmission. These results prompt further investigation into how ABCA7 mutations influence the development and severity of other neurological disorders characterized by altered glutamatergic neurotransmission, such as epilepsy, depression, schizophrenia, Parkinson’s disease, and Huntington’s disease. Significantly, they raise the question of how these mutations might affect the efficacy of treatments for the latter conditions.

### 4.7 Study limitations

While our study provides novel insights into the role of ABCA7 in Glu-GABA imbalance and AD-related neurotoxicity, several limitations should be acknowledged. Our proteomic analyses were performed on bulk whole-brain tissue; it does not provide spatial resolution of region-specific alterations. Thus, subtle differences confined to particular brain regions may have been diluted in the global dataset. Future studies using region-specific or cell-type specific proteomic analyses could provide more detailed insights. We provided interpretation of neurotransmission relies on protein and metabolite levels rather than direct measurements of neuronal activity. Electrophysiological recordings or *in vivo* microdialysis would provide more functional confirmation of the altered glutamatergic and GABAergic signaling suggested by our omics data. Finally, while lipidomic analyses identified lipids associated with ABCA7 deficiency and AD pathology, the mechanistic links between lipid alterations, protein changes, and behavioral phenotypes remain correlative. Further studies are needed to directly establish causal pathways. Taken together, these limitations do not diminish the significance of our findings but rather outline important new avenues for future work to expand upon our observations and strengthen the mechanistic framework of ABCA7 in AD pathology.

## Ethics approval

The animal study and breeding protocols were approved by Forsøksdyrutvalget/KPM or Mattilsynet (FOTS 30157, IV1-2022).

## Consent for publication

Not applicable.

## Data Availability Statement

The mass spectrometry proteomics data have been deposited in the ProteomeXchange Consortium via the PRIDE partner repository (http://www.ebi.ac.uk/pride/archive/):

1. Data of the whole brain proteomics dataset has been published with identifier PXD053250 and the project name “ABCA7 transporter functional role on Alzheimer’s disease onset and progression”.
2. The CSF and ISF dataset has been published with identifier PXD046154 and project name “Alzheimer’s Disease Biomarker Discovery and Validation using Proteomics Analyses of Cerebrospinal Fluid (CSF) and Interstitial Fluid (ISF)”.

The mass spectrometry lipidomics and metabolomics datasets have been deposited via the MassIVE partner repository:

1. with the dataset identifier MSV000094626 (https://doi.org/doi:10.25345/C5HT2GP4W).

## CRediT authorship contribution statement

Anna Maria Górska: writing - original draft, writing - review & editing, investigation, formal analysis, conceptualization, validation, data curation, methodology.

Irene Santos-García: writing - review & editing, investigation, validation, data curation, methodology.

Aleš Kvasnička: writing - review & editing, investigation, validation, data curation, methodology, formal analysis.

Dana Dobešová: writing - review & editing, investigation, validation, data curation, methodology, formal analysis.

David Friedecký: writing - review & editing, validation, supervision, project administration, funding acquisition, resources, data curation, methodology.

Jacob Gildenblat: writing - review & editing, software.

Jens Pahnke: conceptualization, writing - original draft, writing – review & editing, visualization, validation, supervision, software, resources, project administration, methodology, investigation, funding acquisition, formal analysis, data curation.

All the authors have read and agreed to the published version of the manuscript.

## Acknowledgments

The authors thank Thomas Brüning and Ivan Eiriz for their technical support.

Mass spectrometry-based proteomics analyses were performed at the Proteomics Core Facility (Dres. Sachin Singh, Maria Stensland, and Tuula Nyman) at the Department of Immunology, University of Oslo/Oslo University Hospital. This facility is supported by the Core Facilities Program of the South-Eastern Norway Regional Health Authority (HSØ) and NAPI (https://www.napi.uio.no, NFR, Norway; #295910).

Schemes were created using BioRender (https://biorender.com/5c6nzqm).

Writing assistance was applied using ChatGPT Omni

## Abbreviations

Aβ: amyloid-β
AD: Alzheimer’s disease
AMPA: α-amino-3-hydroxy-5-methyl-4-isoxazolepropionic acid
APP: amyloid-β precursor protein
APOE: apolipoprotein E
ABCA7: ATP-binding cassette sub-family A member 7
BBB: blood-brain barrier
CNS: central nervous system
CSF: cerebrospinal fluid
GABA: γ-aminobutyric acid
Glu: glutamate
GFAP: glial fibrillary acidic protein
Gln: glutamine
GWAS: genome-wide association studies
iGluR: ionotropic glutamate receptor
ISF: interstitial fluid
α-KG: α-ketoglutarate
LC-MS/MS: liquid chromatography with tandem mass spectrometry
LV: lateral ventricle
LTP: long-term potentiation
NCSTN: nicastrin
NMDA: N-Methyl-D-Aspartate
OPT: optic tract
PC: Phosphatidylcholine
PCA: Principal component analysis
PD: Parkinson’s disease
PE: Phosphatidylethanolamine
PICK-1: Protein interacting with C-kinase
PLA2: phospholipase A2
PSEN1: presenilin 1
ROS: Reactive oxygen species
TCA: tricarboxylic acid

