## Supplementary Information - figures and tables for "ABCA7 deficiency exacerbates glutamate excitotoxicity in Alzheimer’s disease mice – a new pharmacological target for Glu-related neurotoxicity"

a* Pahnke Lab, www.pahnkelab.eu.

b* Translational Neurodegeneration Research and Neuropathology Lab, Department of Clinical Medicine (KlinMed), Medical Faculty, University of Oslo (UiO), Sognsvannsveien 20, 0372 Oslo, Norway.

c* Section of Neuropathology Research, Department of Pathology, Clinics for Laboratory Medicine (KLM), Oslo University Hospital (OUS), Sognsvannsveien 20, 0372 Oslo, Norway.

d Laboratory for Inherited Metabolic Disorders, Department of Clinical Biochemistry, University Hospital Olomouc and Faculty of Medicine and Dentistry, Palacký University Olomouc, Czech Republic.

e Institute of Nutritional Medicine (INUM), University of Lübeck (UzL) and University Medical Center Schleswig-Holstein (UKSH), Ratzeburger Allee 160, 23538 Lübeck, Germany.

f Department of Neuromedicine and Neuroscience, Faculty of Medicine and Life Sciences, University of Latvia, Jelgavas iela 3, 1004 Riga, Latvia.

g Department of Neurobiology, School of Neuroscience, Biochemistry and Biophysics, The Georg S. Wise Faculty of Life Sciences, Tel Aviv University, Tel Aviv, 6997801, Israel.

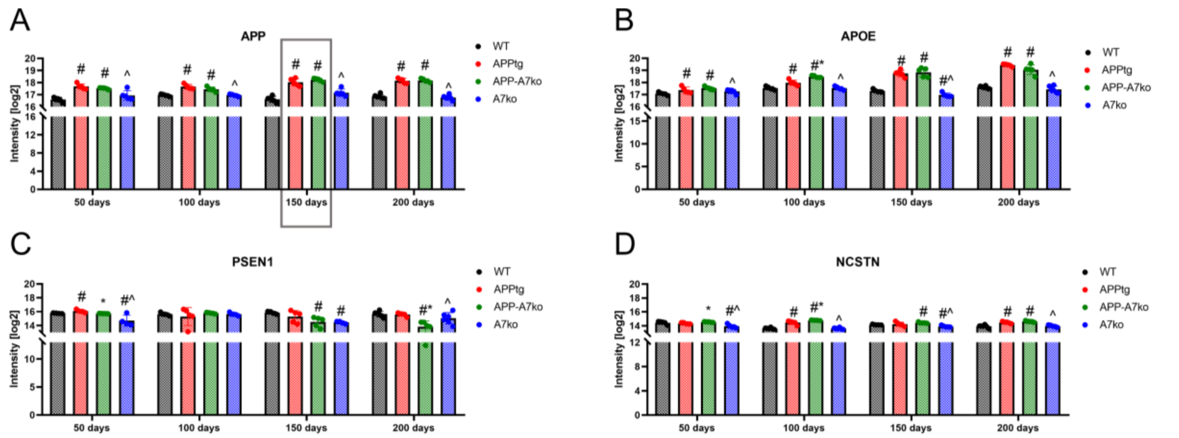


**Fig. S1.** 150 days of age is a crucial time point for the disease progression. Protein intensity values from mass spectrometry measurements of (A) APP, (B) APOE, (C) PSEN1, and (D) NCSTN in the brain of mice aged 50, 100, 150 and 200 days (WT, APPtg, APP-A7ko and A7ko mice). After normality check, statistical analyses were performed with a two-tailed unpaired Student's t-test. $p< 0.05$ was considered to be significant, * vs APPtg; ^ vs APP-A7ko; # vs WT. Data are presented as the means ± SD.

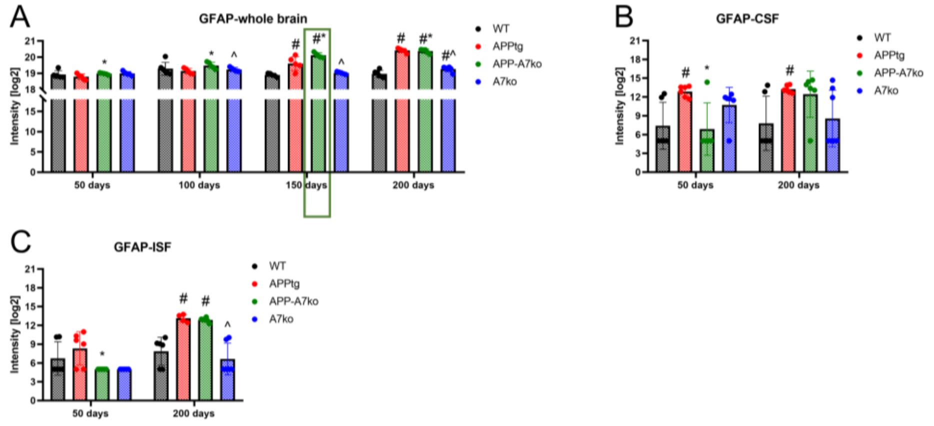


**Fig. S2.** Brain GFAP levels are increased earlier in ABCA7 knockout AD mice. Intensity values from mass spectrometry measurements of GFAP in (A) brain tissue, (B) CSF, and (C) ISF of 50, 100, 150, and 200 days old WT, APPtg, APP-A7ko, and A7ko mice. After a normality check, statistical analyses were performed with a two-tailed unpaired Student's t-test. $p < 0.05$ was considered to be significant, * vs APPtg; ^ vs APP-A7ko; # vs WT. Data are presented as the mean ± SD.

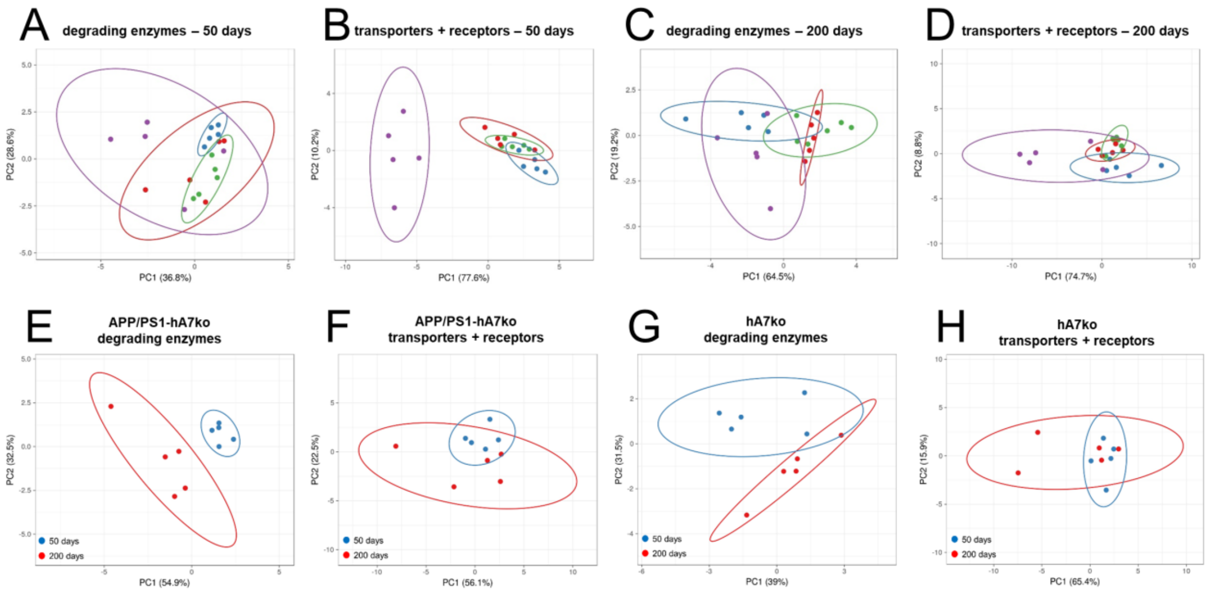


**Fig S3.** (A) PCA for degrading enzymes and (B) transporters plus receptors in 50 days and (C, D) 200-day-old mice, respectively. PCA showing the differences between age (50 days vs. 200 days of age) in the level of degrading enzymes and transporters plus receptors in APP-A7ko (E, F) and A7ko (G, H) mice.

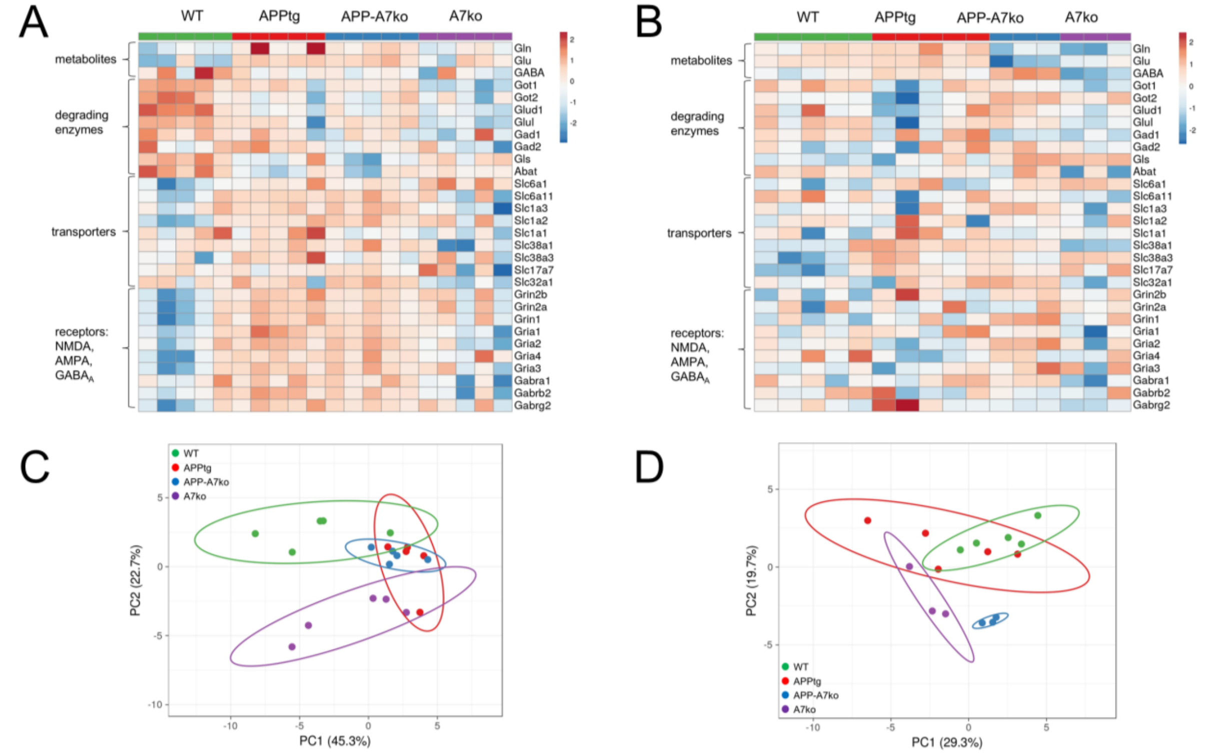


**Fig. S4.** Lack of ABCA7 in AD and control mice affects Glu-GABA-Gln turnover and neurotransmission. Heat map analysis of metabolites (Glu, Gln, GABA) and proteins involved in Glu-GABA-Gln cycle in (A) 100 days and (B) 150 days old control (WT), AD mice (APPtg), AD mice with ABCA7 deficiency (APP-A7ko) and animals without functional

ABCA7 transporters (A7ko). Principal component analysis (PCA) plot showing the percentage of variability between samples due to the genotype effect in (C) 100 and (D) 150-day-old mice.

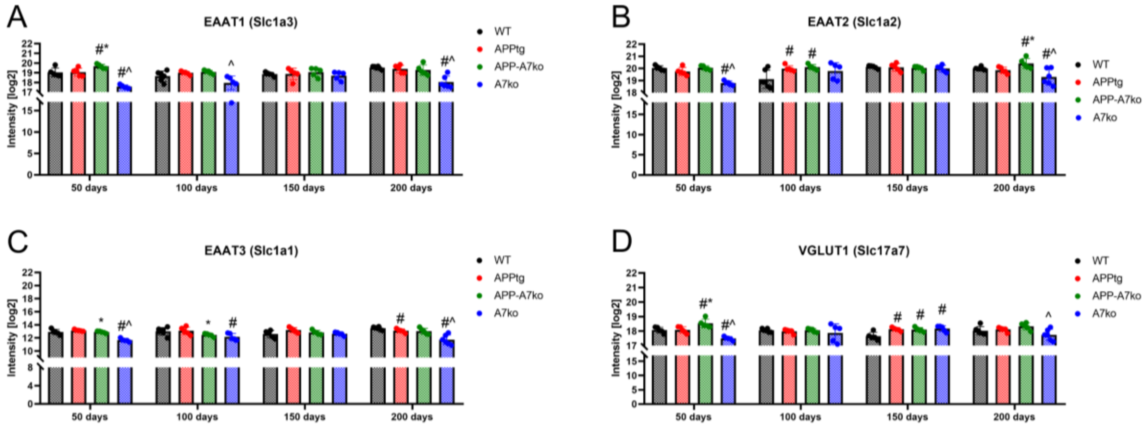


**Fig S5.** The deficit of ABCA7 (A7ko) decreases the level of excitatory amino acid transporters (EAATs) and vesicular glutamate transporter (VGLUT1), having opposite effects in animals with Aβ deposition (APP-A7ko). Intensity values of Glu transporters: (A) EAAT1, (B) EAAT2, (C) EAAT3, and (D) VGLUT1 in the brain of 50, 100, 150, and 200 days old WT, APPtg, APP-A7ko, and A7ko mice. After a normality check, statistical analyses were performed with a two-tailed unpaired Student's t-test. $p < 0.05$ was considered to be significant, * vs APPtg; ^ vs APP-A7ko; # vs WT. Data are presented as the means ± SD.

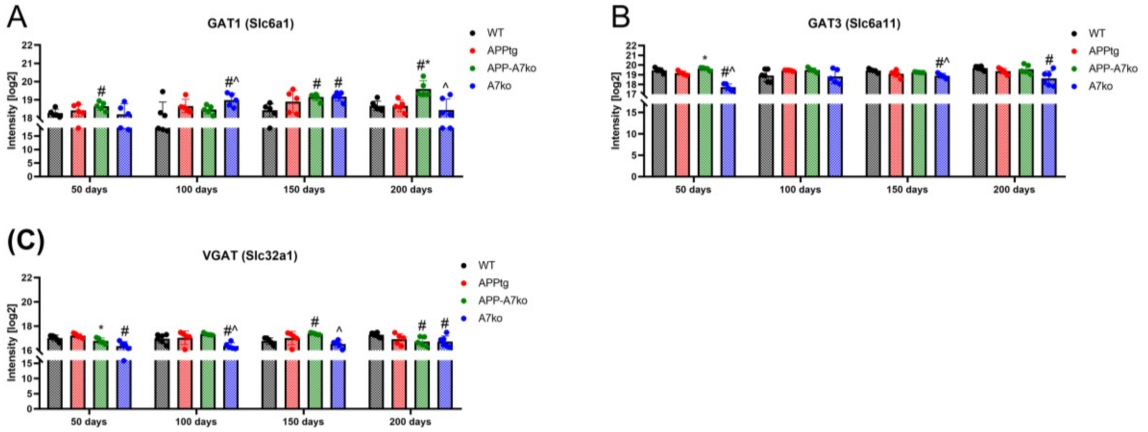


**Fig S6.** Knockout of ABCA7 transporters impaired GABA reuptake. Intensity values of GABA transporters: (A) GAT1, (B) GAT3, and (C) VGAT in the brain of 50, 100, 150 and 200 days old WT, APPtg, APP-A7ko and A7ko mice. Statistical analyses were performed with two-tailed unpaired Student's t-test after normality check. $p < 0.05$ was considered to be significant, * vs APPtg; ^ vs APP-A7ko; # vs WT. Data are presented as the means ± SD.

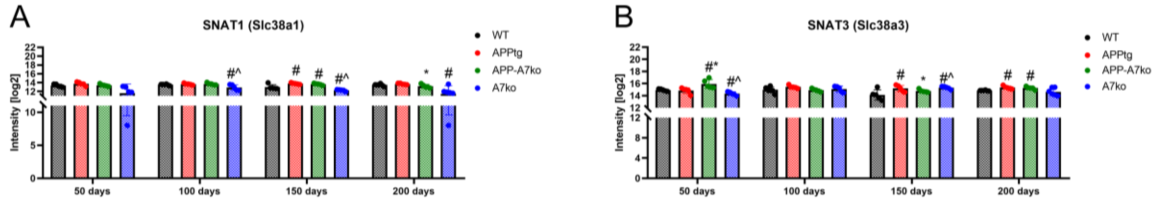


**Fig S7.** ABCA7 deficit impaired Gln delivery from astrocytes to neurons. Intensity values of (A) SNAT1 (neuronal) and (B) SNAT3 (astrocytic) transporters in the brain of 50, 100, 150, and 200 days old WT, APPtg, APP-A7ko, and A7ko mice. After a normality check, statistical analyses were performed with a two-tailed unpaired Student's t-test. $p < 0.05$ was considered to be significant, * vs APPtg; ^ vs APP-A7ko; # vs WT. Data are presented as the means ± SD.

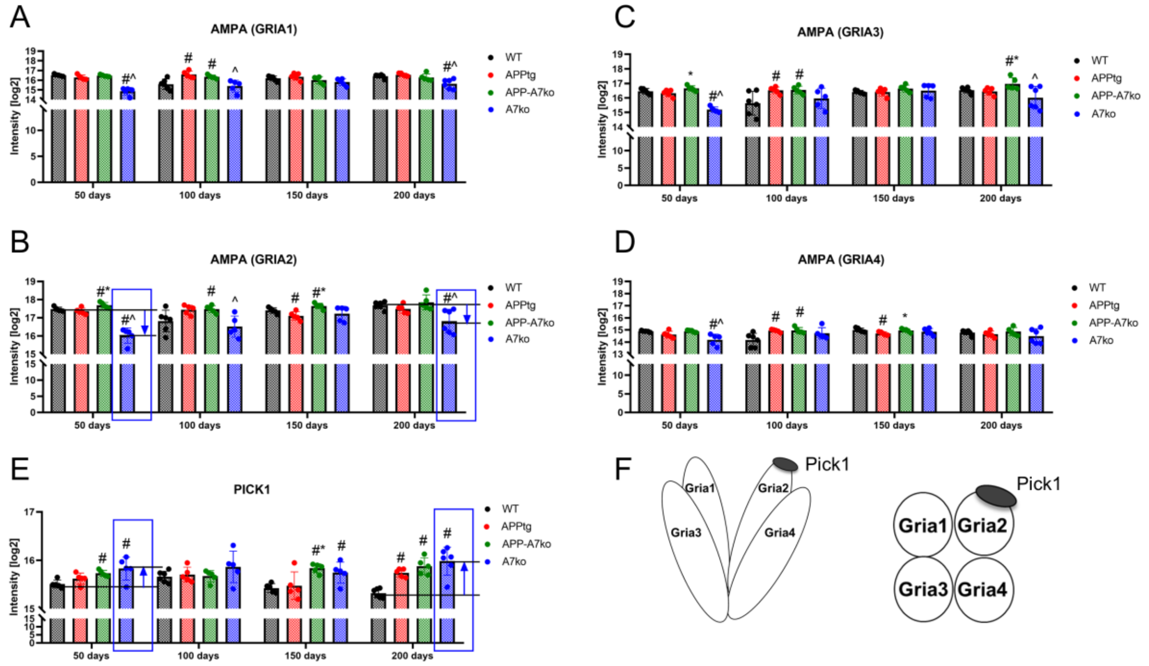


**Fig. S8.** Pick1 induced internalization of AMPA receptors through GRIA2-PICK1 protein-protein interaction in ABCA7 knockout mice. Intensity values of AMPA receptor subunits: (A) GRIA1, (B) GRIA2, (C) GRIA3, (D) GRIA4, and (E) PICK1 in the brain of 50, 100, 150 and 200 days old WT, APPtg, APP-A7ko and A7ko mice. (F) AMPAR subunit composition. After a normality check, statistical analyses were performed with a two-tailed unpaired Student's t-test. $p < 0.05$ was considered to be significant, * vs APPtg; ^ vs APP-A7ko; # vs WT. Data are presented as the means ± SD

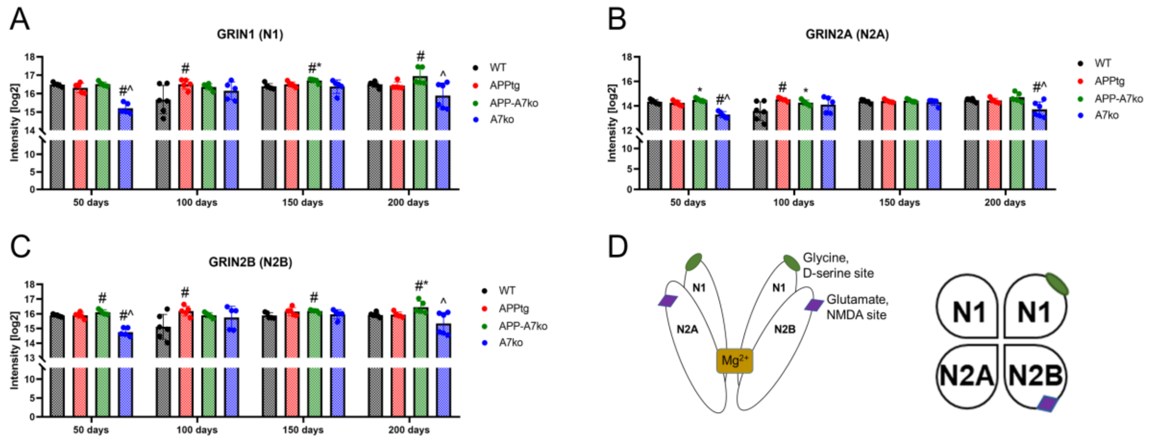


**Fig. S9.** Knockout of ABCA7 transporters decreases the level of NMDAR subunits, having the opposite effect in APPtg animals. Intensity values of NMDA receptor subunits: (A) GRIN1 (N1), (B) GRIN2A (N2A) and (C) GRIN2B (N2B) in the brain of 50, 100, 150 and 200 days old WT, APPtg, APP-A7ko and A7ko mice. (D) NMDAR subunit composition. After a normality check, statistical analyses were performed with two-tailed unpaired Student's t-test. $p < 0.05$ was considered to be significant, * vs APPtg; ^ vs APP-A7ko; # vs WT. Data are presented as the mean ± SD.

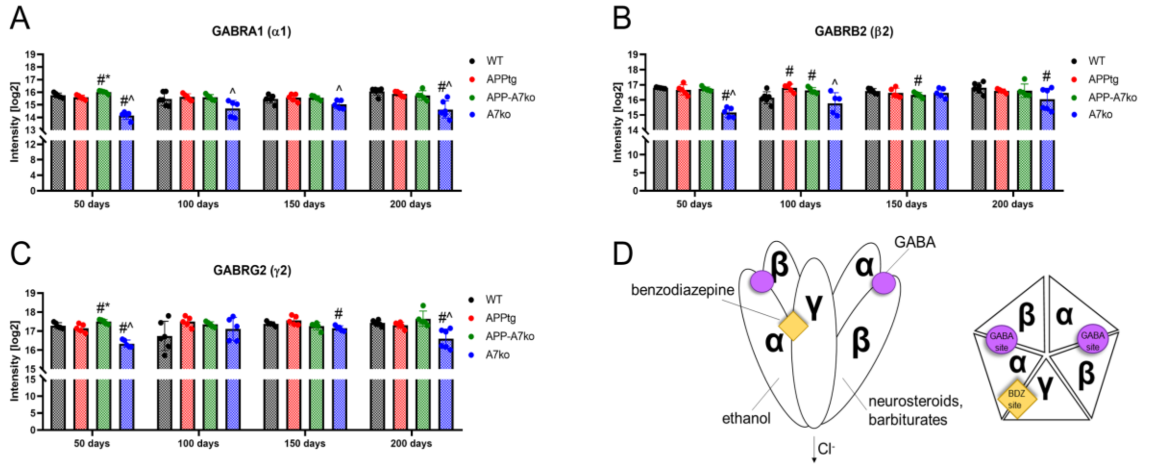


**Fig. S10.** Lack of ABCA7 decreases the level of all $GABA_A$ receptor subunits. Intensity values of $GABA_A$ receptor subunits: (A) GABRA1 (α1), (B) GABRB2 (β2) and (C) GABRG2 (γ2) in the brain of 50, 100, 150 and 200 days old WT, APPtg, APP-A7ko and A7ko mice. (D) $GABA_A$ subunit composition. After a normality check, statistical analyses were performed with a two-tailed unpaired Student's t-test. $p < 0.05$ was considered to be significant, * vs APPtg; ^ vs APP-A7ko; # vs WT. Data are presented as the mean ± SD.

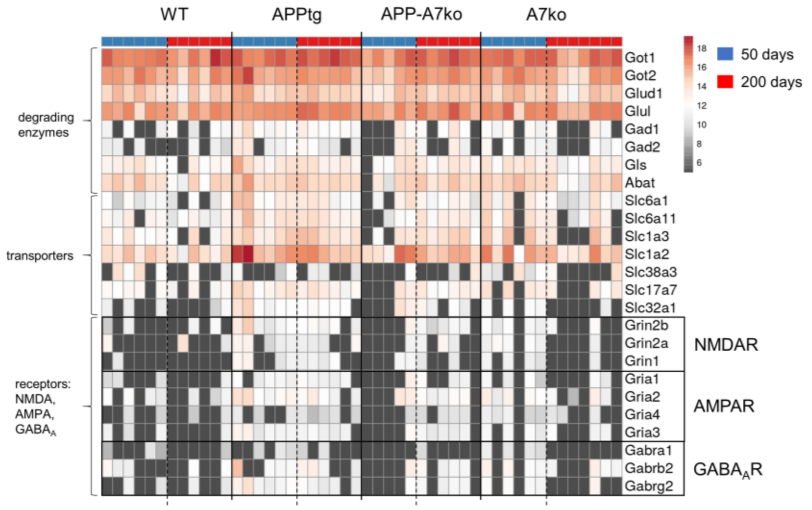


**Fig. S11.** Aβ deposition in the brain increases the level of NMDA, AMPA, and GABA$_A$ receptors subunits in CSF. Heat map analysis of proteins involved in Glu-GABA-Gln turnover and neurotransmission in CSF of 50 days and 200 days old control (WT), AD (APPtg) mice with (APP-A7ko) ABCA7 transporters deficiency and animals without functional ABCA7 transporters (A7ko). Dark grey squares represent no expression of the protein in this animal.

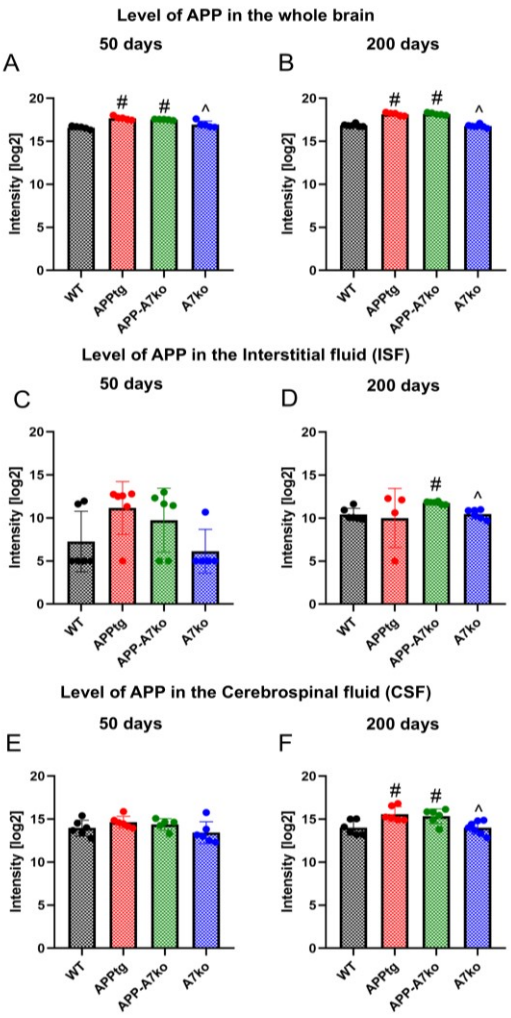


**Fig. S12.** Progressive deposition of Aβ in the brain induced infiltration of APP into ISF and CSF. Intensity values of APP in brain tissue (A, B), ISF (C, D), and CSF (E, F) of 50 and 200-day-old WT, APPtg, APP-A7ko, and A7ko mice. After a normality check, statistical analyses were performed with two-tailed unpaired Student's t-test. $p < 0.05$ was considered to be significant, * vs APPtg; ^ vs APP-A7ko; # vs WT. Data are presented as the mean ± SD.

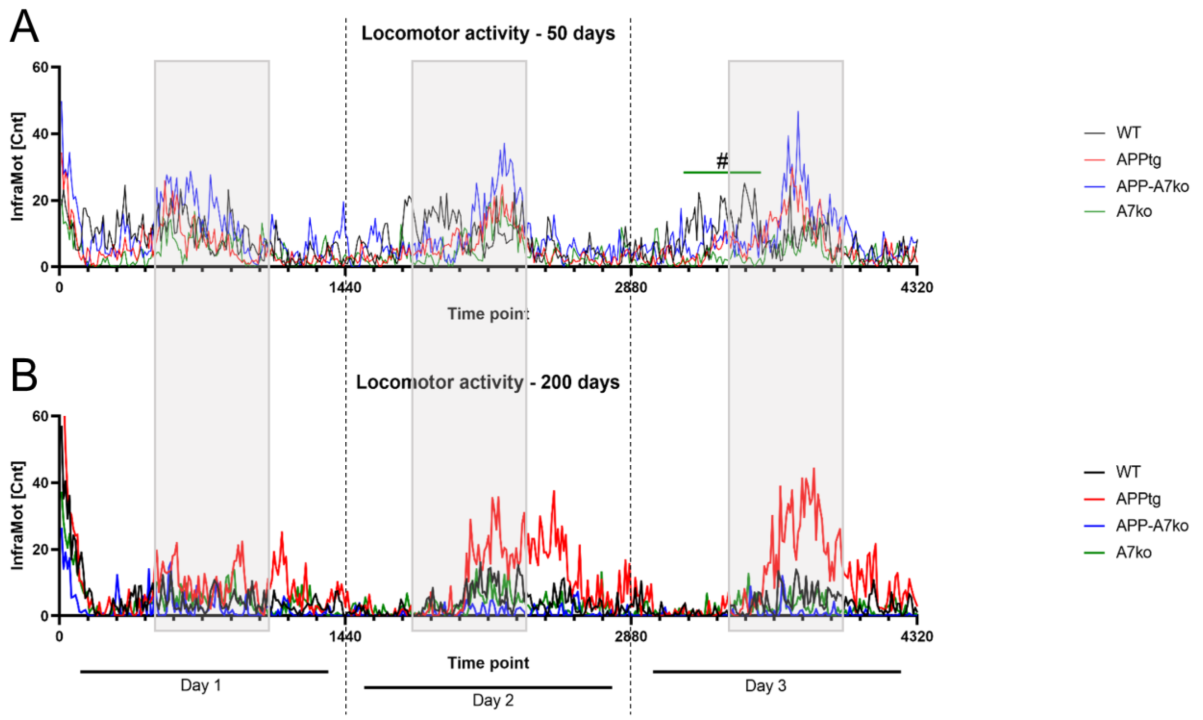


**Fig. S13.** Lack of ABCA7 decreases locomotor activity in control and AD mice. Spontaneous activity of mice of (A) 50 days and (B) 200 days was measured every 10 minutes for 72h, each day for 1440 minutes. The pattern of locomotor activity of animals from each group is presented as a mean value. Statistical analyses were performed using a two-tailed unpaired Student's t-test after a normality check. $p < 0.05$ was considered to be significant, # A7ko vs WT.

**Fig. S14.** List of all analyzed lipids according to the order presented on Heat maps.

FA: FA(14:0), FA(15:0), FA(16:0), FA(16:1), FA(18:0), FA(18:1), FA(18:2), FA(18:3), FA(20:1), FA(20:2), FA(20:3), FA(20:4), FA(22:4), FA(22:5), FA(22:6)

PG: PG(32:0), PG(16:0/16:0), PG(32:1), PG(34:1), PG(16:0/18:1), PG(18:0/18:1), PG(18:1/18:1) , PG(18:0/20:4), PG(38:4)

PI: PI(32:0), PI(16:0/16:0), PI(32:1), PI(16:0/16:1), PI(34:0), PI(16:0/18:0), PI(16:0/18:1), PI(18:0/16:1), PI(34:1), PI(16:0/18:2), PI(18:1/16:1), PI(34:2), PI(18:0/18:1), PI(36:1), PI(18:0/18:2), PI(18:1/18:1), PI(36:2), PI(16:0/20:3), PI(18:1/18:2), PI(36:3), PI(16:0/20:4), PI(36:4), PI(37:4), PI(38:3), PI(18:0/20:3), PI(16:0/22:4), PI(18:0/20:4), PI(38:4), PI(16:0/22:5), PI(18:1/20:4), PI(38:5), PI(16:0/22:6), PI(38:6), PI(18:0/22:4), PI(40:4), PI(18:0/22:6), PI(40:6)

PS: PS(16:0/18:1), PS(34:1), PS(18:0/18:1), PS(36:1), PS(36:2), PS(18:0/20:4), PS(38:4)

Cholesterol

Cer: Cer(d18:1/16:0), Cer(d18:1/18:0), Cer(d18:1/18:1), Cer(d18:1/22:4), Cer(d18:1/22:5), Cer(d18:1/22:6), Cer(d18:2/22:5)

Hex2Cer: Hex2Cer(d16:1/16:0), Hex2Cer(d18:1/14:0), Hex2Cer(d18:1/16:0)

LPC: LPC(14:0), LPC(15:0), LPC(16:0), LPC(16:1), LPC(17:0), LPC(17:1), LPC(18:0), LPC(18:1), LPC(18:2), LPC(18:3), LPC(20:0), LPC(20:1), LPC(20:2), LPC(20:3), LPC(20:4), LPC(22:0), LPC(22:1), LPC(22:4), LPC(22:5), LPC(24:0), LPC(25:0), LPC(26:0), LPC(26:1), LPC(O-16:0), LPC(P-16:0), LPC(O-18:0), LPC(P-18:0), LPC(O-18:2), LPC(O-18:3)

LPE: LPE(16:1), LPE(18:2), LPE(20:3), LPE(22:6)

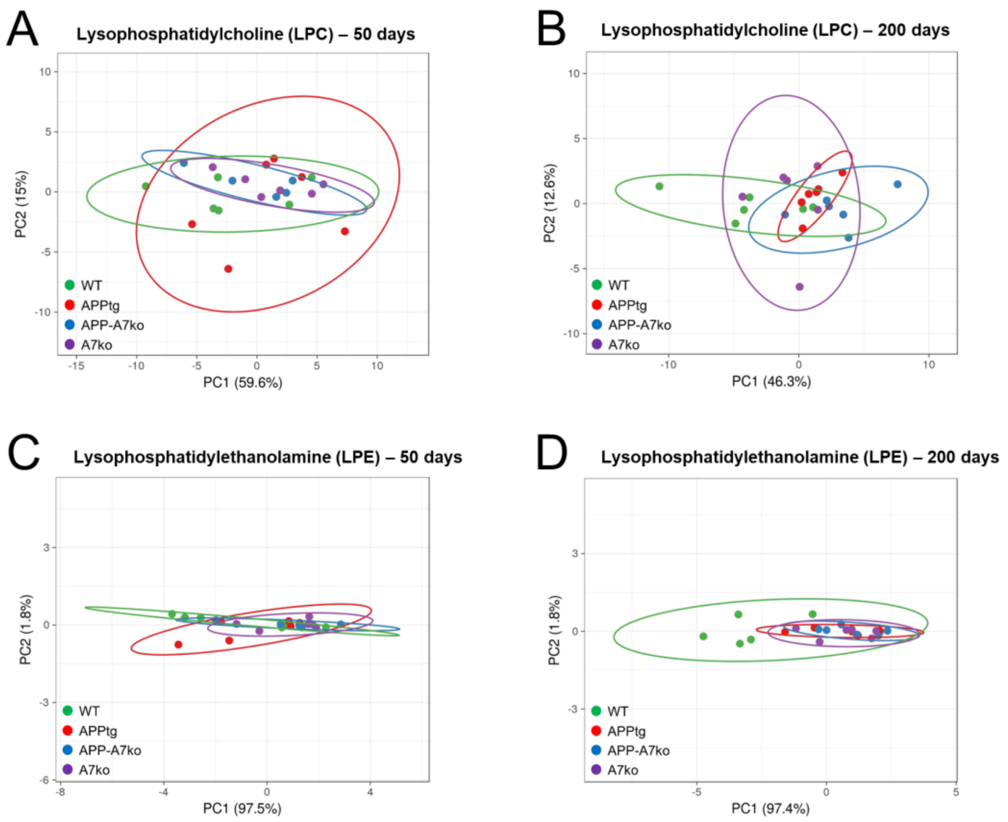


**Fig. S15.** The principal component analysis (PCA) plot shows the percentage of variability in (A, B) LPC and (C, D) LPE levels between samples due to genotype effects in mice that are 50 and 200 days of age, respectively.

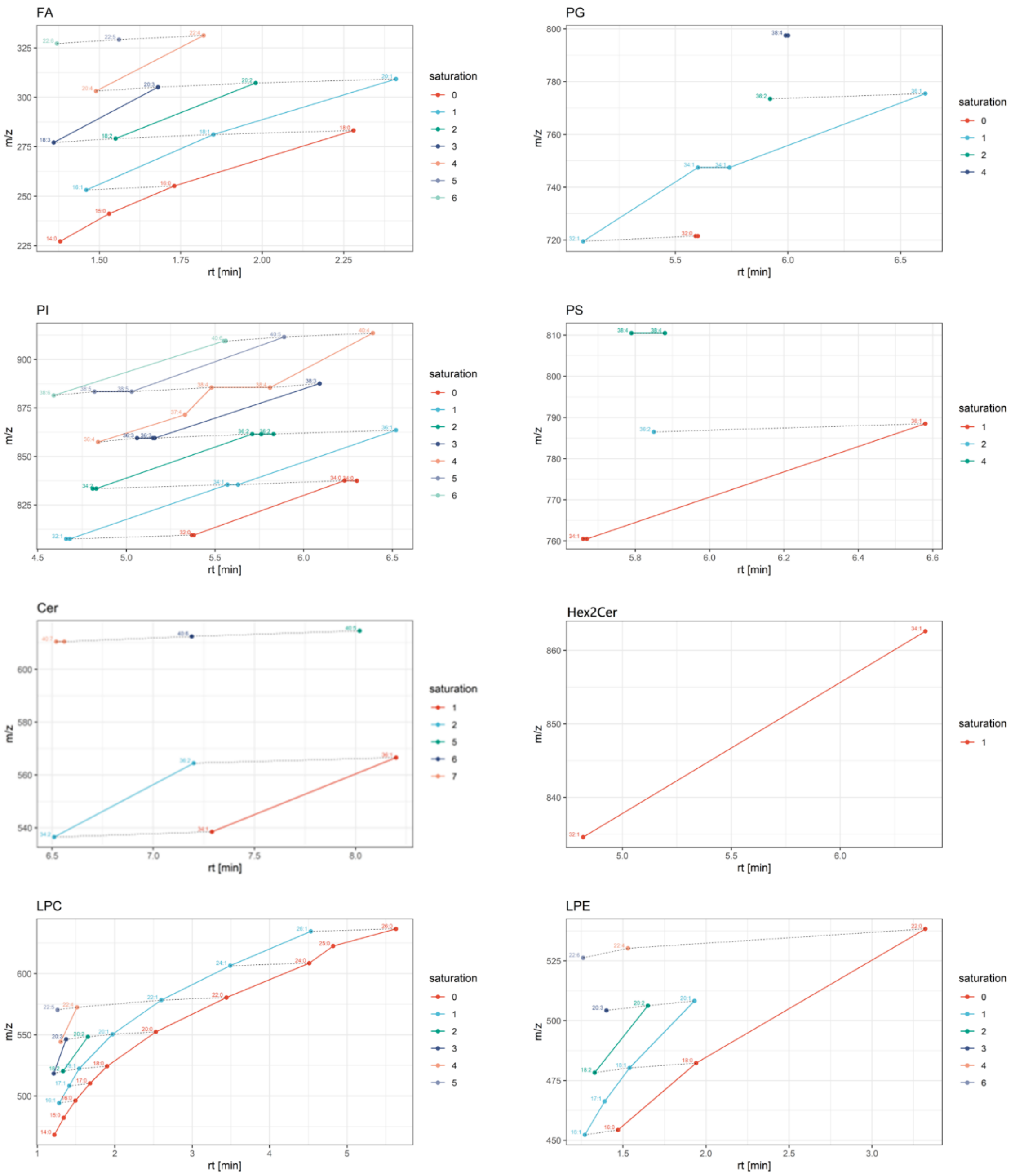


**Figure S16.** Retention behavior of lipids plotted as mass-to-charge ratio (m/z) versus retention time (in min). The effect of both the acyl chain length (number of carbons) and the degree of unsaturation (number of double bonds) on retention time was used to verify correct lipid identification and annotation.
